# Specificity and sensitivity of an RNA targeting type III CRISPR complex coupled with a NucC endonuclease effector

**DOI:** 10.1101/2021.09.13.460032

**Authors:** Sabine Grüschow, Catherine S. Adamson, Malcolm F. White

## Abstract

Type III CRISPR systems detect invading RNA, resulting in the activation of the enzymatic Cas10 subunit. The Cas10 cyclase domain generates cyclic oligoadenylate (cOA) second messenger molecules, activating a variety of effector nucleases that degrade nucleic acids to provide immunity. The prophage-encoded *Vibrio metoecus* type III-B (VmeCmr) locus is uncharacterised, lacks the HD nuclease domain in Cas10 and encodes a NucC DNA nuclease effector that is also found associated with Cyclic-oligonucleotide-based anti-phage signalling systems (CBASS). Here we demonstrate that VmeCmr is activated by target RNA binding, generating cyclic-triadenylate (cA_3_) to stimulate a robust NucC-mediated DNase activity. The specificity of VmeCmr is probed, revealing the importance of specific nucleotide positions in segment 1 of the RNA duplex and the protospacer flanking sequence (PFS). We harness this programmable system to demonstrate the potential for a highly specific and sensitive assay for detection of the SARS-CoV-2 virus RNA with a limit of detection (LoD) of 2 fM using a commercial plate reader without any extrinsic amplification step. The sensitivity is highly dependent on the guide RNA used, suggesting that target RNA secondary structure plays an important role that may also be relevant *in vivo*.

## INTRODUCTION

Early bioinformatic analyses of CRISPR associated (*cas*) genes revealed the existence of a “Repeat associated mysterious protein” (RAMP) operon that included a gene encoding a predicted polymerase (1). Subsequently, a seminal study of Cas protein families classified the RAMP (Cmr) and Mtube (Csm) subtypes, each including a putative polymerase gene (*cmr*2 and *csm*1, respectively) (2). The burgeoning CRISPR nomenclature, which had become increasingly confusing for everyone, was (first) standardised in 2011, resulting in the classification of the polymerase protein as Cas10 – the signature protein of Type III CRISPR systems (3).

The first biochemical studies resulted in the observation that type III CRISPR systems comprise a multi-subunit ribonucleoprotein complex that uses CRISPR RNA (crRNA) to target and degrade cognate RNA molecules (4). In a follow up study, the Terns and Li labs determined the crystal structure of Cas10, confirming the predicted PALM polymerase/cyclase domain and ruling out a role for the N-terminal HD nuclease domain in the targeted degradation of RNA (5). Target RNA cleavage was subsequently shown to be catalysed by the Cas7 (Cmr4/Csm3) subunit, which forms the backbone of Type III complexes (6–9). This left the function of the Cas10 subunit unresolved – an uncertainty that was compounded by the observation that plasmid clearance by a type III system from *Sulfolobus islandicus* required transcription of DNA target genes and depended on the presence of an uncharacterised Cas protein known as Csx1 (10). Similarly, plasmid clearance in *Staphylococcus epidermidis* was shown to be dependent on the active site of the polymerase/cyclase domain and the presence of the Csm6 protein (11). Csx1 and Csm6 are found adjacent to many Type III CRISPR systems, although not part of the multi-protein complex. Tellingly, both proteins have a C-terminal HEPN nuclease domain (12), suggesting a possible function as ribonucleases for these proteins. *In vivo* studies demonstrated that the *S. epidermidis* type III system tolerated lysogenic phage but destroyed lytic phage via transcription-dependent DNA targeting (13).

The first convincing evidence that the HD domain of Cas10 was a nuclease came from a combined structure / function study of the protein from *Thermococcus onnurineus*, which demonstrated that the HD nuclease domain has ssDNA endonuclease activity *in vitro* (14).This was confirmed unequivocally by studies of intact type III complexes from *Streptococcus thermophilus*, *Pyrococcus furiosus*, *Thermotoga maritima*, *S. islandicus and Thermus thermophilus* which demonstrated ssDNA-specific HD nuclease activity, activated by target RNA binding (15–19).This DNase activity is not ubiquitous however, with some type III systems having little detectable RNA-activated DNase activity, at least *in vitro* (20,21).

In 2017, another major piece of the puzzle was resolved with the demonstration that the PALM domains of Cas10 generate a novel family of cyclic oligoadenylate (cOA) molecules once activated by target RNA binding (22,23). cOA, which is generated in a range of ring sizes from cA_3_ to cA_6_, functions as a second messenger of infection, activating effector nucleases to provide immunity against mobile genetic elements. These effectors include the widespread Csx1/Csm6 family, which bind cA_4_ or cA_6_ in their CARF (CRISPR associated Rossman fold) domains, resulting in activation of the HEPN ribonuclease domain at the C-terminus (22–27) and extensive non-specific degradation of RNA in the cell (28). Subsequently, CARF-family effector proteins capable of targeting DNA when activated by cOA have also been described (29–31). All of these enzymes are presumed to carry out “collateral cleavage” by degrading the nucleic acids of both host and virus, delaying viral replication. This process can lead to cell death if not controlled, but a range of specialised ring nuclease enzymes that degrade cOA allow cells to deactivate the collateral cleavage activity and survive viral assault (reviewed in (32)).

A subset of Cas10 orthologues, primarily found in type III-B CRISPR systems in the gamma proteobacteria, firmicutes and bacteroidetes, lack the N-terminal HD domain entirely and are thus smaller proteins, around 600 aa in length (33). The proteins tend to have the cyclase active site residues conserved in the PALM domains, suggesting that they provide interference solely via cyclic nucleotide signalling rather than direct Cas10-mediated DNA degradation. Here we focus on the system from *Vibrio metoecus*, a type III-B system comprising subunits Cmr1-6 where the Cmr2/Cas10 subunit lacks an HD-nuclease domain, together with a Cas6 enzyme for crRNA generation (Figure 1A). The CRISPR system is encoded on a prophage that is also found in several *Vibrio cholerae* strains, and there are no associated adaptation genes (34). Adjacent to the *cmr* genes is a gene encoding NucC, a hexameric DNA endonuclease that was recently shown to be activated by cyclic trinucleotides in cyclic-oligonucleotide-based anti-phage signalling systems (CBASS) (35). NucC is thus an interesting example of an effector protein found in both CBASS and CRISPR defence systems. In the latter context it is assumed to be activated by cA_3_ generated by the activated Cas10 subunit, although this has not been demonstrated directly.

**Figure 1.**
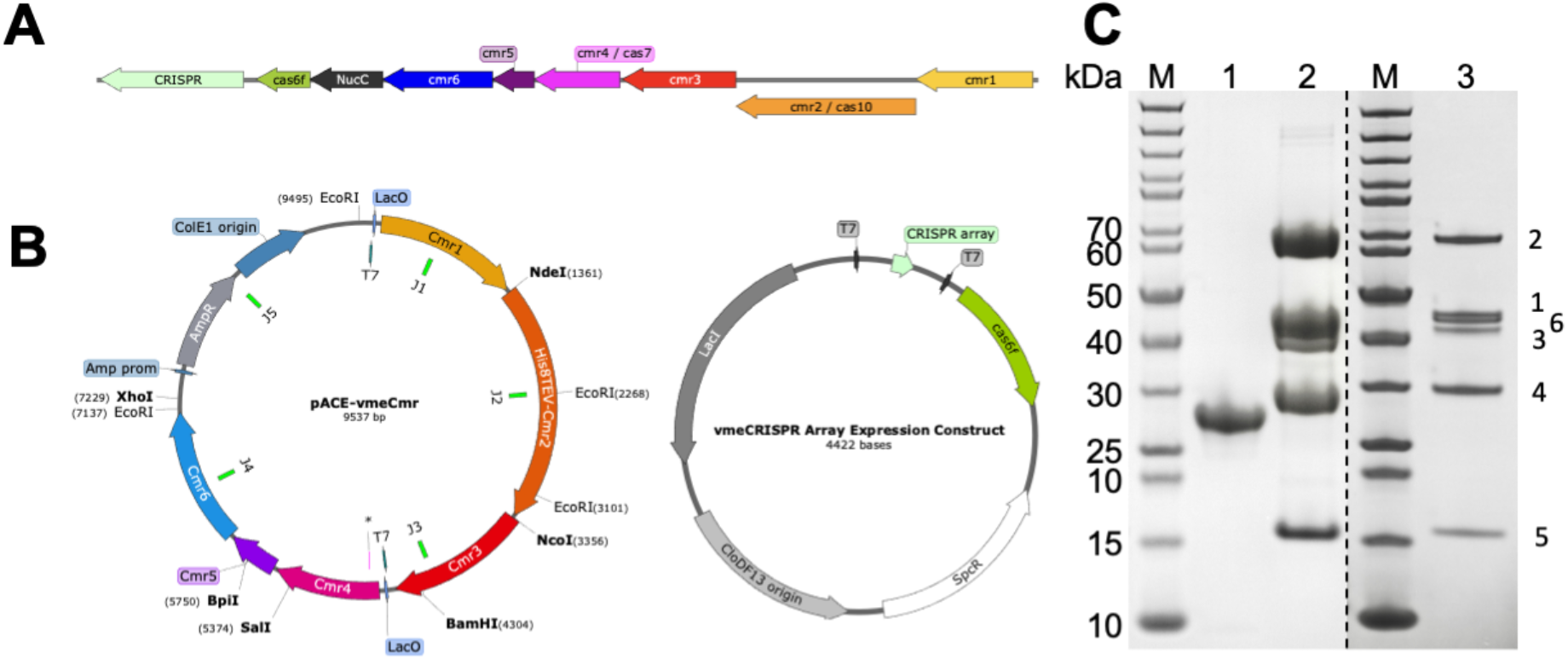
Plasmid expression construct and SDS-PAGE of purified proteins. **A**: Genomic locus for the type III-B CRISPR locus of *Vibrio metoecus*. **B**: Expression construct pACE-vmeCmr for the VmeCmr complex, with the junctions (J1-J5) for Gibson assembly indicated, and the expression construct for the CRISPR array and Cas6 protein. Gene locus and plasmid maps were generated with SnapGene Viewer. **C**: SDS-PAGE analysis of the purified recombinant proteins. M: PageRuler Unstained (Thermo Scientific); lane 1: NucC; lane 2: VmeCmr wild type complex; lane 3: VmeCmr complex loaded at lower concentration to allow visualisation of all subunits. The dashed line separates two different gels. The VmeCmr subunit number is shown to the right of the gel.

In the past few years, the collateral cleavage activities of Cas12 and Cas13 have been repurposed for the development of rapid and specific assays capable of detecting specific RNA or DNA sequences (36–39). Two recent studies have demonstrated that the ability of type III (Cas10) CRISPR systems to detect RNA specifically and synthesise cOA can be harnessed in combination with CARF family effector nuclease Csm6 to provide an alternative assay system (40,41). Here, we demonstrate that a prophage-encoded *Vibrio* type III-B system generates cA_3_ on specific target RNA binding, resulting in the activation of the NucC effector nuclease for non-specific dsDNA degradation. By coupling these activities to a fluorescent readout, we obtain a limit of detection of 2 fM for SARS-CoV-2 RNA targets.

## MATERIALS AND METHODS

### Cloning

Enzymes were purchased from Thermo Scientific or New England Biolabs and used according to manufacturer’s instructions. Oligonucleotides and synthetic genes were obtained from Integrated DNA Technologies (IDT, Coralville, Iowa, USA). Synthetic genes were codon-optimized for *E. coli* and restriction sites for cloning incorporated where necessary. All final constructs were verified by sequencing (GATC Biotech, Eurofins Genomics, DE). *Vibrio metoecus nucC* (NucC) was obtained as a G-Block with flanking restriction sites for cloning. After digestion with NcoI and SalI, *nucC* was ligated into NcoI and XhoI-digested pEV5HisTEV (42) to allow expression as a N-terminal His8-fusion protein.

The expression construct pACE-Cmr for the *V. metoecus* Cmr interference complex (VmeCmr) was assembled using the NEBuilder^®^ HiFi DNA Assembly kit (New England Biolabs) following the Gibson assembly strategy (43). All fragments were PCR-amplified from synthetic genes prior to assembly. The final construct (Figure 1B) contained a ColE1 origin of replication and ampicillin resistance gene, copied from pACE (MultiColi^™^, Geneva Biotech, Genève, CH). The expression of *cmr1-3* and *cmr4-6* was driven by T7 promoters. The *cmr2* (*cas10*) gene included a sequence encoding a TEV-cleavable, N-terminal His_8_-tag.

The *V. metoecus* CRISPR array was designed as two repeat sequences flanking two oppositely directed BpiI recognition sites (5’-gt<u>gtcttcg</u>tacctt<u>gaagac</u>ca) to allow later insertion of the target/spacer sequence of choice. The synthetic cassette also contained flanking NcoI and SalI sites that were used to clone the pre-array into MCS-1 of pCDFDuet^™^-1 (Novagen, Merck Millipore). *Vibrio metoecus cas6f*, obtained as a G-Block with flanking NdeI and XhoI sites, was subsequently cloned into MCS-2 of the latter construct to give pCDF-notarget_CRISPR (Figure 1B). The VmeCas6 protein is 69 % identical to the Cas6f protein from a *Schewanella putrfaciens* type I-F system, and the CRISPR sequence is closely related, suggesting a 5’-handle sequence 5’-CUUAGAAA (44). It has been suggested that a *Vibrio* phage has co-opted this hybrid type III-B/I-F system for inter-phage conflict (34).

Target/spacer sequences were obtained as synthetic oligonucleotides with a 5’-overhang sequence of 5’-GAAA for the sense strand and 5’-GAAC for the antisense strand. After the two strands were annealed, they were ligated into BpiI-digested pCDF-notarget_CRISPR to give pCDF-*target*_CRISPR. vmeRepeat and spacer sequences are listed in Table 1.

**Table 1.**
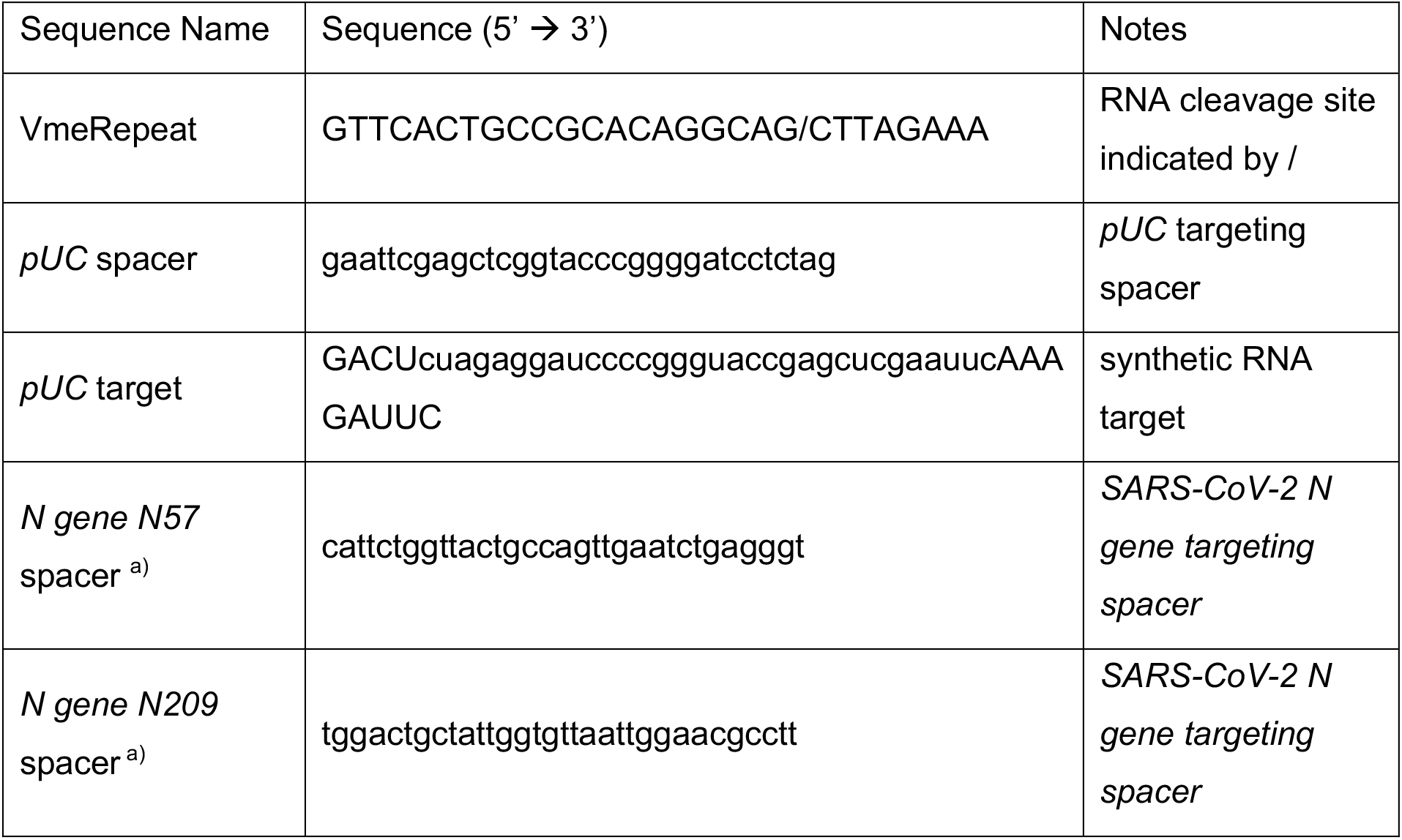
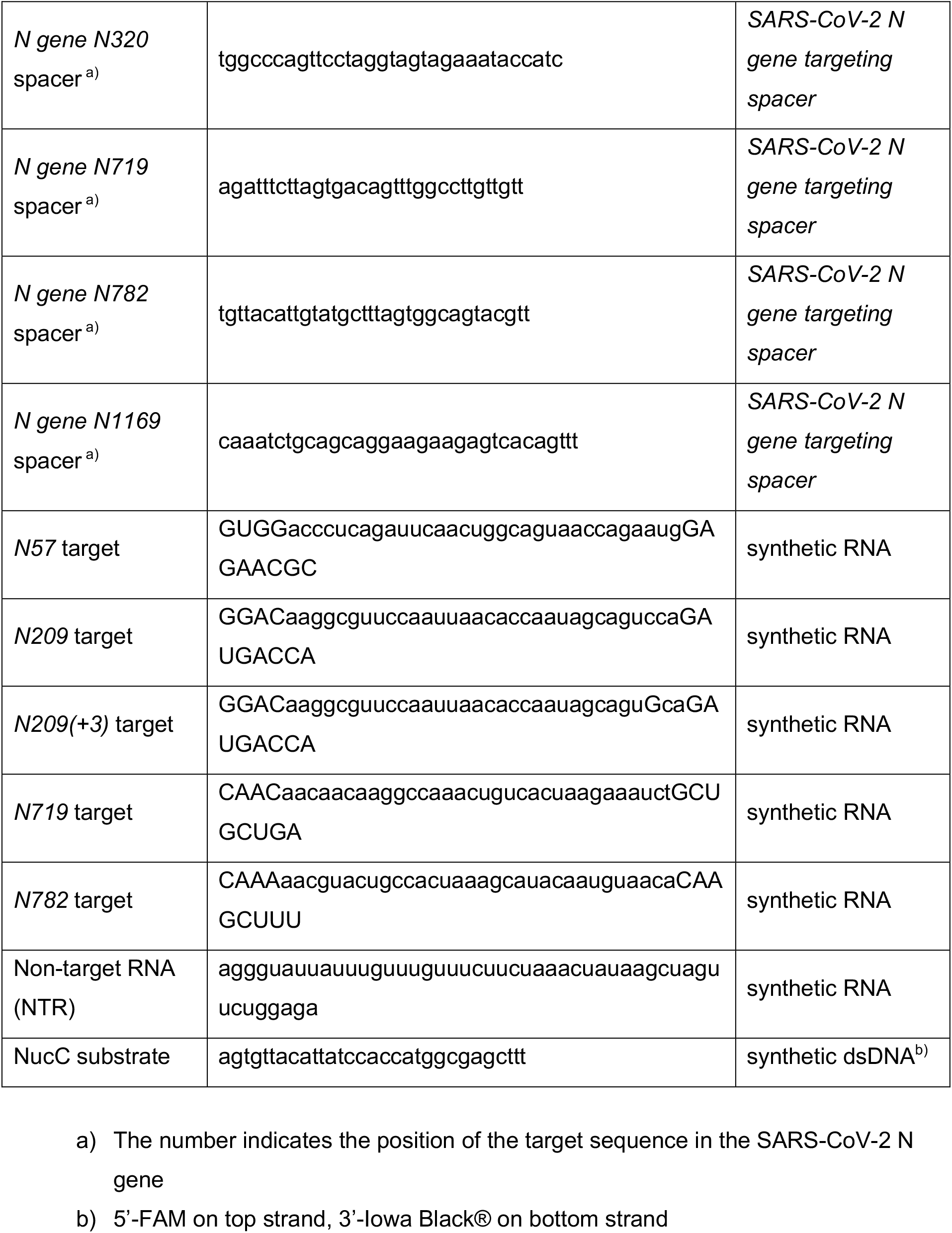

### RNA targets and NucC reporter substrate

RNA oligonucleotides were obtained from IDT. The mRNA for the SARS-CoV-2 N gene was obtained by *in* vitro transcription as follows. The template for *in vitro* transcription was constructed by PCR-amplification (primers: 5’-ctagccatgGCGTTTGAGACGGGCGACAG and 5’-ctaggtcgaCAGCTCGCTGGTCCAGAACTG) from the commercially available plasmid 2019-nCoV_N_Positive Control (IDT) and cloned by restriction digest / ligation into MCS-1 of pRSFDuet^™^-1 (Novagen) to give pSG220. The N gene plus the upstream T7 promoter region was PCR-amplified from pSG220 using Phusion^™^ High-Fidelity DNA polymerase (Thermo Scientific) and standard primers ACYCDuetUP1 and DuetDOWN1. The agarose gel purified PCR product served as template for *in vitro* transcription using the MEGAscript^®^ T7 kit (Invitrogen) according to the manufacturer’s protocol. After DNase I digest the transcript was purified by phenol:chloroform extraction / *iso*-propanol precipitation. The integrity of the transcript was verified by agarose gel electrophoresis. RNA species were quantified by spectrophotometry using a dilution series for each sample; for RNA oligomers the molar concentration was obtained from OD_260_ measurement and the calculated extinctions coefficients (IDT); N gene transcript concentration was estimated using the generic ε_260_ of 0.025 (ng μl^−1^)^−1^ cm^−1^ and a molecular weight of 464,564 Da (from the formula: MW = 1,449 nt * 320.5 Da + 159 Da). All RNA species were dispensed into single use aliquots and stored at −70 °C.

The NucC reporter substrate was obtained by heating and slow cooling of the 5’-FAM- and 3’-Iowa Black^®^-labelled strands (25 μM each) in 10 mM Tris-HCl, pH 8, 50 mM NaCl. The dsDNA substrate was stored in the dark at −20 °C.

### SARS-CoV-2 Genomic RNA Extraction

All infectious work with human coronavirus SARS-CoV-2 (strain hCOV-19/England/2/2020; kind gift of Dr Marian Killip, Public Health England, UK) was conducted using a class II Microbiology safety cabinet inside a Biosafety Level 3 (BSL3) biocontainment facility. Vero E6 cells (African green monkey kidney epithelial cell, ECACC, 85020206) were routinely cultured in Dulbecco Modified Eagle Medium (DMEM) supplemented with 10% v/v fetal bovine serum (FBS) and used to generate a high-titer SARS-CoV-2 working stock. The virus stock was generated by infecting Vero E6 cells at a multiplicity of infection (MOI) of 0.01 in DMEM supplemented with 1 % v/v FBS. Cell supernatant was harvested 72 hours post-infection, clarified by centrifugation for 5 min at 276 × g, aliquoted, flash frozen in liquid nitrogen and stored at −80 °C. Virus stock titer was determined by plaque assay using a confluent monolayer of Vero E6 cells in 12-well plates, infected with 10-fold serial dilutions of virus stock (200 μL/well) and incubated for 1 hour with shaking at 37 °C. The virus inoculum was removed, and cells overlaid with 1 % v/v avicel (Dupont) in DMEM supplemented with 2 % FBS and incubated at 37 °C / 5% CO_2_ for 72h. Cells were then fixed with 5 % v/v paraformaldehyde/phosphate buffered saline (PBS) and plaques visualized by crystal violet staining (0.2 % w/v). The virus titer was determined to be 6 × 10^6^ plaque forming units (PFU)/mL. Ten-fold serial dilution of the SARS-CoV-2 stock were prepared using PBS and viral genomic RNA extracted using the QIAamp viral RNA mini extraction kit (Qiagen) according to the manufacturers spin protocol and the recommended 5.6 μg polyA carrier RNA/140 μL virus sample. The QIAamp viral RNA extraction kit procedure inactivates SARS-CoV-2 (45) therefore the eluted RNA was removed from the BSL3 lab and stored as single-use aliquots at −70 °C until required for analysis. The extracts were diluted two-fold in RNase-free water just before use for RT-qPCR analysis and for the VmeCmr/NucC coupled assay.

### Protein Production and Purification

*E. coli* BL21 Star^™^ (DE3) cells (Invitrogen) were co-transformed with pACE-Cmr and pCDF-*target*_CRISPR. The wild type VmeCmr construct and a Cmr4-D26A variant designed to abolish target RNA cleavage were expressed and purified in the same way. Overnight cultures were diluted 100-fold into LB containing 100 μg ml^−1^ ampicillin and 50 μg ml^−1^ spectinomycin, incubated with shaking at 37 °C, 180 rpm until the OD_600_ reached 0.8. After induction with 200 μM IPTG, incubation was continued at 27 °C overnight. Cells were harvested by centrifugation and pellets stored at −20 °C. Cells were resuspended in lysis buffer (50 mM Tris-HCl, 500 mM NaCl, 20 mM imidazole, 10 % (v/v) glycerol, pH 7.5) and lysed by sonication. The cleared lysate was loaded onto a pre-equilibrated HisTrap Crude FF (GE Healthcare) column, washed with lysis buffer and eluted in a gradient with increasing imidazole concentration (to 0.5 M). VmeCmr complex-containing fractions were pooled, dialysed at 4 °C overnight in the presence TEV protease against lysis buffer without imidazole. The protein solution was passed through the HisTrap Crude FF column a second time, the VmeCmr complex-containing flow-through was concentrated using an Amicon Ultracentrifugal filter (30 kDa MWCO, Merck-Millipore) and further purified by size exclusion chromatography (HiPrep^™^ 16/60 Sephacryl^®^ pg 300 HR, Cytiva) using 20 mM Tris-HCl, 500 mM NaCl, 10 % (v/v) glycerol, pH 7.5 as mobile phase. Fractions from the main peak (Vol_Ret_ 53 ml) containing the VmeCmr complex were combined, concentrated as above and the NaCl concentration adjusted to 50 mM by dilution with gel filtration buffer lacking NaCl before further purification over a HiTrap Heparin column (Cytiva). The sample was loaded in 20 mM Tris-HCl, 500 mM NaCl, 10 % (v/v) glycerol, pH 7.5 and eluted with a NaCl gradient. Fractions containing the VmeCmr complex, eluting at approximately 200 mM NaCl, were combined and concentrated as before to 3 – 4 mg ml^−1^, dispensed into single-use aliquots and flash-frozen in liquid nitrogen. Aliquots were stored at −70 °C. Protein concentrations were determined by UV quantitation (NanoDrop 2000, Thermo Scientific) using calculated extinction coefficients (ExPASy, ProtParam software for protein; AAT Bioquest for crRNA).The concentration of VmeCmr complex was estimated using an extinction coefficient of 610,240 M^−1^ cm^−1^, which was obtained by adding the values for the protein component in Cmr1_1_ Cmr2_1_ Cmr3_1_ Cmr4_4_ Cmr5_3_ Cmr6_1_ stoichiometry and an estimated value for the crRNA (calculated 396,900 M^−1^ cm^−1^ at 260 nm, estimated 200,000 M^−1^ cm^−1^ for 280 nm).

The production and purification of NucC was carried out as for the VmeCmr complex with the following exceptions: *E. coli* C43(DE3) was used as the expression host, size exclusion chromatography was carried out on a HiLoad^®^ 16/60 Superdex^®^ pg 200 column (GE Healthcare) using 20 mM Tris-HCl, 250 mM NaCl, 10 % (v/v) glycerol, pH 7.5 as mobile phase, and no heparin purification was performed. The NucC peaks eluting at approximately 160 ml and 180 ml during gel filtration were collected separately, and likely correspond to hexameric and trimeric species, respectively, as observed previously (35). The trimeric species was used for this study. Quantitation was carried out as described for the VmeCmr complex using a calculated extinction coefficient of 29,910 M^−1^ cm^−1^, and concentrations are reported for the NucC trimer. SDS-PAGE analysis of all proteins used in this study is shown in Figure S1.

### Liquid chromatography of cyclic oligonucleotides

To determine which cOAs are produced by VmeCmr, 500 nM VmeCmr [*pUC*_CRISPR] and 5 μM *pUC* target were incubated at 37 °C in 12.5 mM Tris-HCl, pH 8.0, 25 mM NaCl, 10 mM MgCl_2_, 10 % (v/v) glycerol, 500 μM ATP for 90 min. Enzyme was removed by spin filtration (Nanosep 3 kDa MWCO, Pall), and the filtrate was analysed by LC-MS. Compounds were separated on a Kinetex^®^ EVO C18 column (2.6 μm, 2.1 × 50 mm, Phenomenex) using a Dionex UltiMate 3000 chromatography system with the following gradient of acetonitrile (B) against 10 mM ammonium bicarbonate (A): 0 – 2 min 2% B, 2 – 10 min 2 – 8% B, 10 – 11 min 8 – 95% B, 11 – 13 min 95% B, 13 – 14 min 95 – 2% B, 14 – 18 min 2% B at a flow rate of 300 μl min^−1^ and column temperature of 40 °C. Elution was monitored by UV at 254 nm and by mass spectrometry (LCQ Fleet mass spectrometer, Fisher Scientific; +ve ion mode, *m/z* 200 - 1800). Data were analysed using Xcalibur^™^ (Thermo Scientific) and visualised in Prism (GraphPad).

To test for degradation of cA_3_ by NucC, 500 nM NucC and 100 μM cA_3_ were incubated at 37 °C for 1 h under the same conditions as for cOA production. Samples were processed and analysed as described above, however, elution was monitored by UV spectrophotometry alone.

### Nuclease assay

All assays were performed in duplicate unless stated otherwise on a FluoStar Omega plate reader (BMG Labtech) using fluorescence detection (l_ex / em_ 485 / 520 nm) in black, non-binding half-area 96-well plates (Corning). Key experiments were performed in at least two independent experiments; representative examples are shown, and data are provided in Supporting Information. Screens for selected assay conditions for both NucC alone and in combination with VmeCmr are also found in Supporting Information. The standard nuclease assay contained 25 – 250 nM wild-type or Cmr4-D26A variant VmeCmr complex in 12.5 mM Tris-HCl, pH 8.0, 10 – 20 mM NaCl, 10 mM MgCl2, 10 % (v/v) glycerol, 500 μM ATP, 125 nM FAM : Iowa Black^®^ double-stranded DNA substrate (Table 1) and varying concentrations of target or control RNA unless stated otherwise. Synthetic target RNAs are listed in Table 1. The reaction was incubated at 37 °C in the absence of NucC to allow for cyclic oligoadenylates to be produced; NucC was added at *t* = 10 min to start the nuclease reaction. Fluorescence was measured throughout in 30 s intervals at 37 °C for up to 50 min. The signal between 2 and 9.5 min was used for baseline subtraction (Mars software, BMG Labtech).

### RT-qPCR

RT-qPCR was performed using the 2019-nCoV RUO Kit (Integrated DNA Technologies, #10006713) in combination with the iTaq^™^ Universal SYBR^®^ Green One-Step Kit (BioRad). Each 20 μl reaction contained 10 μl 2X iTaq universal SYBR^®^ Green reaction mix, 0.25 μl iScript reverse transcriptase, 2 μl premixed N2 primer/probe mix, and 2 μl sample. The reverse transcriptase was omitted to test for DNA contamination. Samples were either a 10-fold serial dilution of SARS-CoV-2 N gene transcript (final concentrations 1 aM – 1 pM) or SARS-CoV-2 extracts. Each condition, including a non-template control, was run in triplicate. The RT-qPCR reaction was performed in a Stratagene Mx5000P instrument using the 2-step protocol for SYBR Green with Dissociation Curves settings and monitoring ROX as reference dye. Cycle conditions were: 10 min at 50 °C; 1 min at 95 °C; 40x 10 s at 95 °C, 30 s at 60 °C; the dissociation curve was measured at 55 – 95 °C. Amplification plots and dissociation curves are shown in Figure S8, Ct values are provided in Table S1.

### Data analysis

Microsoft Excel and Prism 8 (Graphpad) were used for data analysis. Statistical analyses were performed with Prism 8 (GraphPad). Target specificity was analyzed by extracting the time point (T_t_) when the fluorescence intensity crossed a threshold value. The threshold value was set to 1/8^th^ of the maximal measured fluorescence signal of the reference substrate (*pUC* target, Table 1). Individual T_t_ values were subtracted from the T_t_ value obtained in the absence of RNA to give ΔT_t_ values. Finally, ΔT_t_ values were normalized to the ΔT_t_ value of the reference substrate.

To determine the limit of detection, the mean of fluorescence intensities between 29 – 31 min reaction time was calculated for each target concentration and independent experiment. Sample means higher than the mean of the signal for the reference reaction (without RNA or with non-target RNA) plus 10 standard deviations were deemed detected. The lowest measured concentration detected was taken as the LoD. The quoted LoDs represent the average LoD from at least two independent experiments.

## RESULTS

### Cloning, expression and purification

In *V. metoecus* NucC (NucC hereafter) is the only effector protein encoded by the CRISPR locus (Figure 1A). To investigate the activity of the VmeCmr / NucC system, we built two plasmid constructs (Figure 1B), one expressing synthetic versions of the codon optimised *cmr*1-6 genes and a second encoding Cas6 and a mini-CRISPR array (see materials and methods). This system can be programmed to detect any RNA sequence by changing the spacer sequence in the CRISPR array. The VmeCas6 enzyme processes the pre-crRNA transcript to generate mature guide RNAs with the 5’-handle CUUAGAAA (44). The Cas6 enzyme is not part of type III effectors and as expected VmeCas6 did not co-purify with VmeCmr. Initially we programmed the system using a single spacer that matched a synthetic target RNA species used previously (21). The VmeCmr complex expressed at a high level in *E. coli* and could be purified in milligram quantities. We cloned, expressed and purified NucC separately, using expression plasmid pEV5HisTEV (42) in *E. coli*, to obtain milligram quantities of pure protein (Figure 1C).

### VmeCmr generates predominantly cA3 when activated by target RNA

We proceeded to characterise the *V. metoecus* type III-B (VmeCmr) system by using target RNA complementary to the crRNA spacer sequence of the ribonucleoprotein complex (see Table 1 for sequences) to activate the cyclase activity of Cas10. We confirmed that VmeCmr generates predominantly cA3 when activated by target RNA, with much lower levels of cA4 made, as expected for a system coupled to NucC (Figure 2).

**Figure 2.**
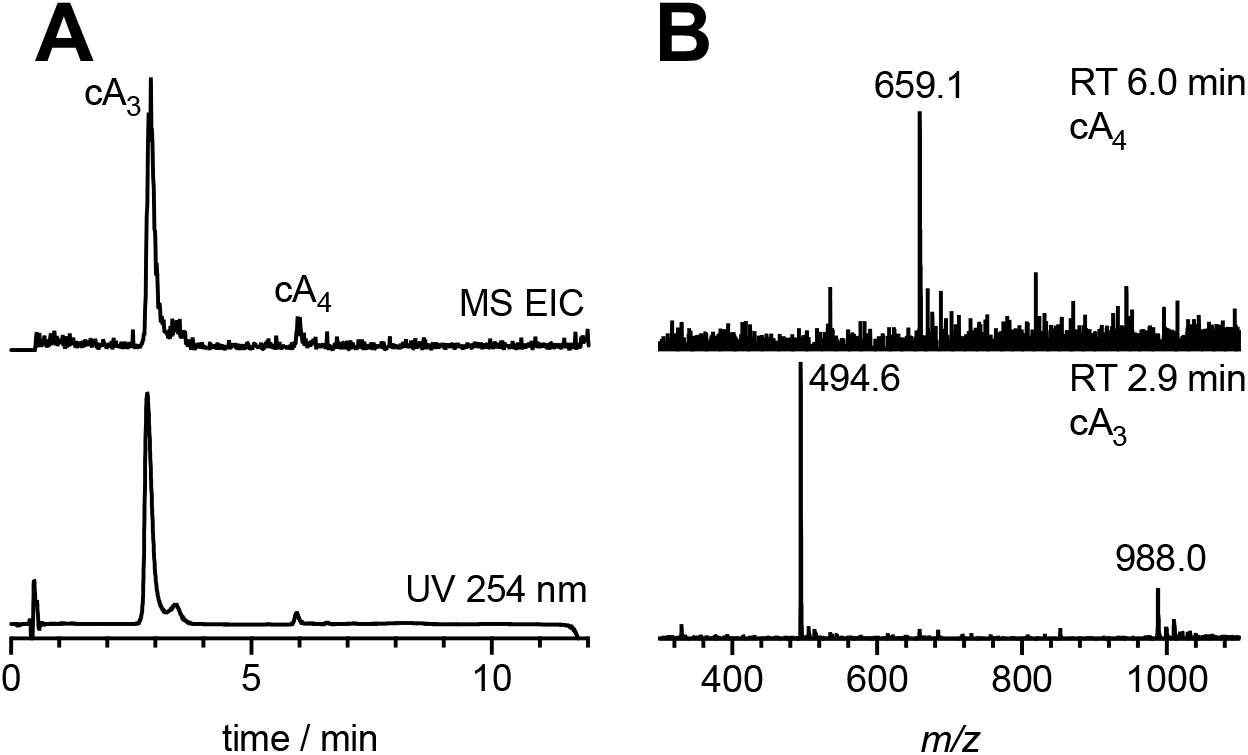
VmeCmr generates predominantly cA3 when activated by target RNA. **A**: LC-MS chromatograms with mass and UV monitoring, respectively. The extracted ion chromatogram (EIC) shows all ion species corresponding to cyclic oligoadenylates cA_2_ – cA_6_. **B**: Mass spectra for products with retention time (RT) 2.9 min and 6.0 min corresponding to cA_3_ and cA_4_. Expected for cA_3_ C_30_H_38_N_15_O_18_P_3_^2+^ *m/z* 494.7, found 494.6; expected for cA_4_ C_40_H_48_N_20_O_24_P_4_^2+^ *m/z* 659.1, found 659.1.

### NucC functions as a cA3 activated nuclease with high sensitivity

NucC was previously shown to cleave plasmid DNA when activated by synthetic cA_3_ (35). To aid development of a continuous assay for NucC, we used a synthetic DNA duplex of 30 bp with a fluorescein reporter dye quenched by IOWA Black^®^ on the opposite strand and monitored the fluorescent signal generated by NucC when activated by its activator cA_3_. We investigated the limit of detection (LoD) of the fluorescent signal generated by 250 nM NucC (trimer concentration) using synthetic cA_3_ in the absence of VmeCmr. The signal reached saturation in less than 5 min for cA_3_ concentrations of ≥ 10 nM, with cA3 concentrations of ≥ 100 nM achieving maximal activation of the system (Figure 3A). We chose the fluorescent signals between 29 – 31 min as our diagnostic signal for subsequent experiments as it provided a good compromise of sensitivity and experiment time. At 30 min, approximately 100 pM cA_3_ activator provided half-maximal nuclease activation (Figure 3C), with 10 pM cA_3_ providing a statistically significant signal after 30 min incubation (Figure 3B), suggesting that NucC has a very high affinity for cA_3_. As the cOA level generated by Cas10 is directly proportional to the RNA present in the sample (26,46), the affinity of effector nucleases for their cOA activators confers intrinsic limitations on the sensitivity of the system.

**Figure 3.**
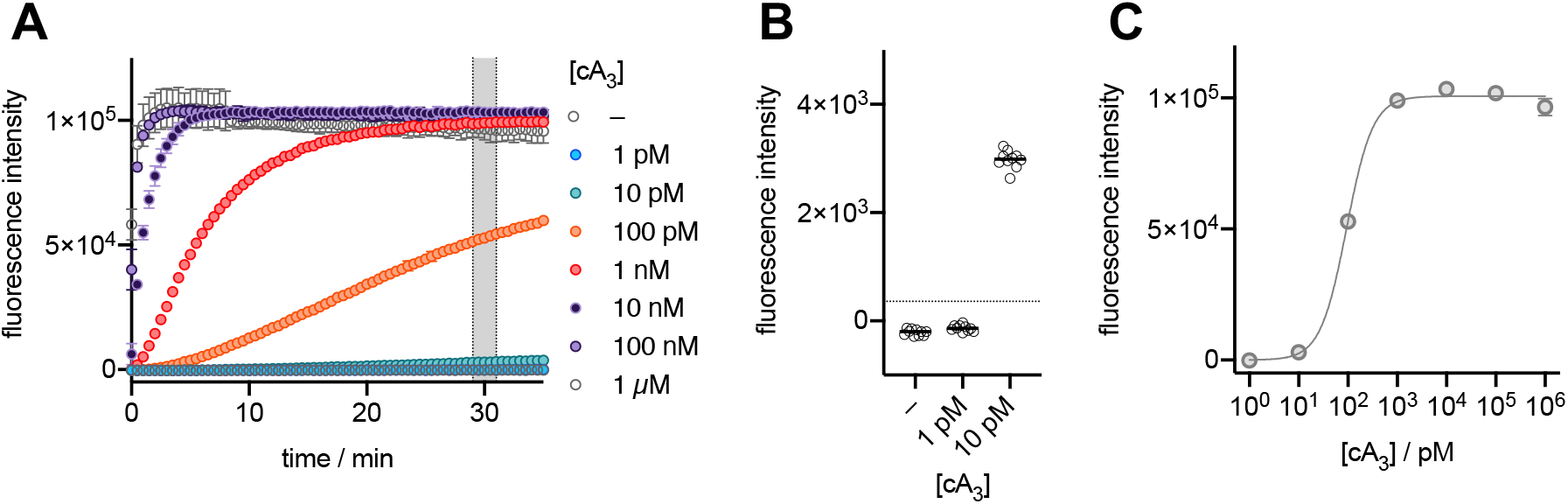
NucC activity using synthetic cA3 as activator. **A**: Fluorescence signal curves at various cA_3_ concentrations. **B**: Fluorescence intensities between 29 – 31 min plotted against cA_3_ concentration. The dotted line represents the threshold value. Samples with fluorescnce intensities higher than the threshold were deemed to activate NucC. The lowest activating cA3 concentration was 10 pM. **C**: The fluorescence intensities between 29 – 31 min reaction time were used for non-linear regression (specific binding with Hill slope) to estimate the cA_3_ concentration at which NucC is 50 % activated (Act50). Act50 was 94 ± 4 pM cA_3_ with a Hill slope of 1.6.

Some Csx1/Csm6 family proteins have an intrinsic “ring nuclease” activity for the degradation of their cOA activator (25,47,48) – an activity that is probably important *in vivo* for the control of the CRISPR mediated immune response (46). To determine whether NucC degrades cA_3_, we incubated 100 μM cA_3_ with 0.5 μM NucC for 60 min and monitored the products by liquid chromatography and UV detection (Figure S2D) after removal of the enzyme by spin filtration (MWCO 3 kDa). The assay showed no significant depletion of cA_3_ and no trace of any degradation products, suggesting that NucC does not degrade its own activator at significant levels.

### Coupling of NucC activation with target RNA detection

We next coupled NucC to specific RNA detection by VmeCmr, monitoring NucC-mediated fluorescent readout as before. We first titrated target RNA concentrations to establish the limits of RNA detection using the NucC-coupled fluorogenic assay (Scheme 1) and a standard reference RNA target sequence (pUC target, Table 1). For wild-type VmeCmr, RNA target concentrations at the sub-picomolar level produced a fluorescent signal within 30 min of initiating the assay that was statistically significant by comparison with a control reaction using 50 nM non-target RNA (Figure 4). RNA target concentrations in the mid-picomolar range, produced a very fast signal that reached its maximum within 10 min.

**Figure 4.**
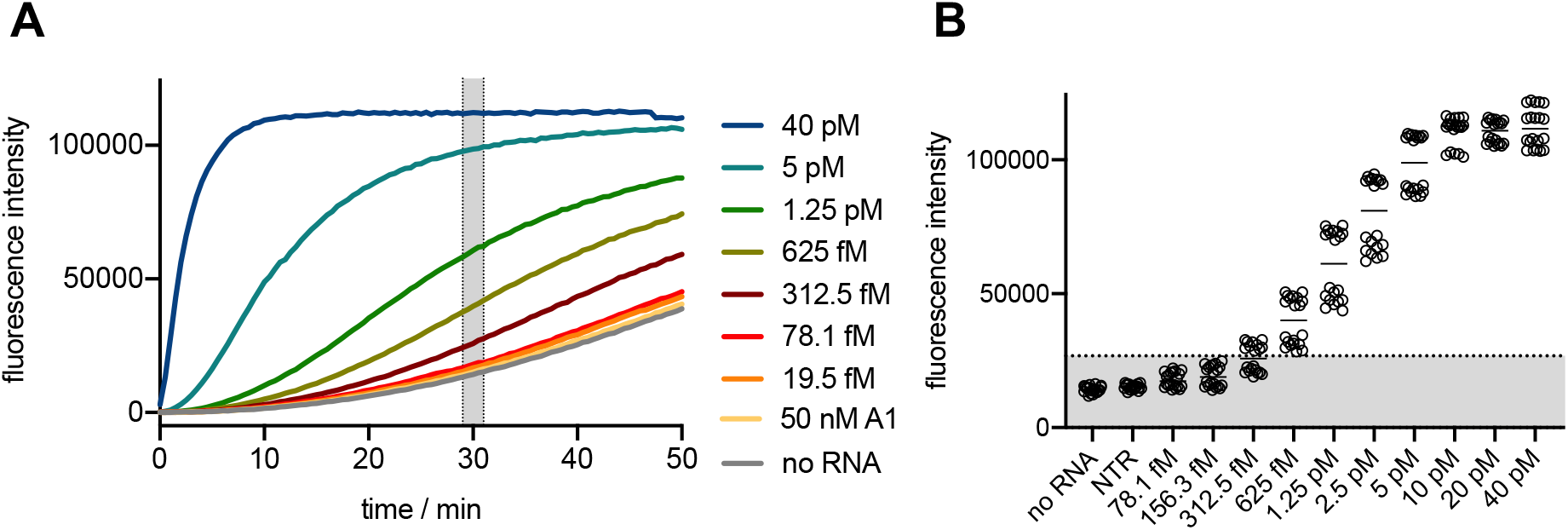
LoD of target RNA for the coupled assay. 250 nM VmeCmr with *pUC*-targeting crRNA was used under standard conditions. A: Fluorescence signal curves in the presence or absence of target RNA. Target RNA concentrations are indicated. The mean of two independent experiments is shown for each RNA concentration. B: The fluorescence intensities of measurements from 29 – 31 min reaction time (shaded in A) were used to determine the LoD. The dotted line represents the detection threshold value which was defined as 10 SDs above the reference (no RNA) mean.

**Scheme 1.**
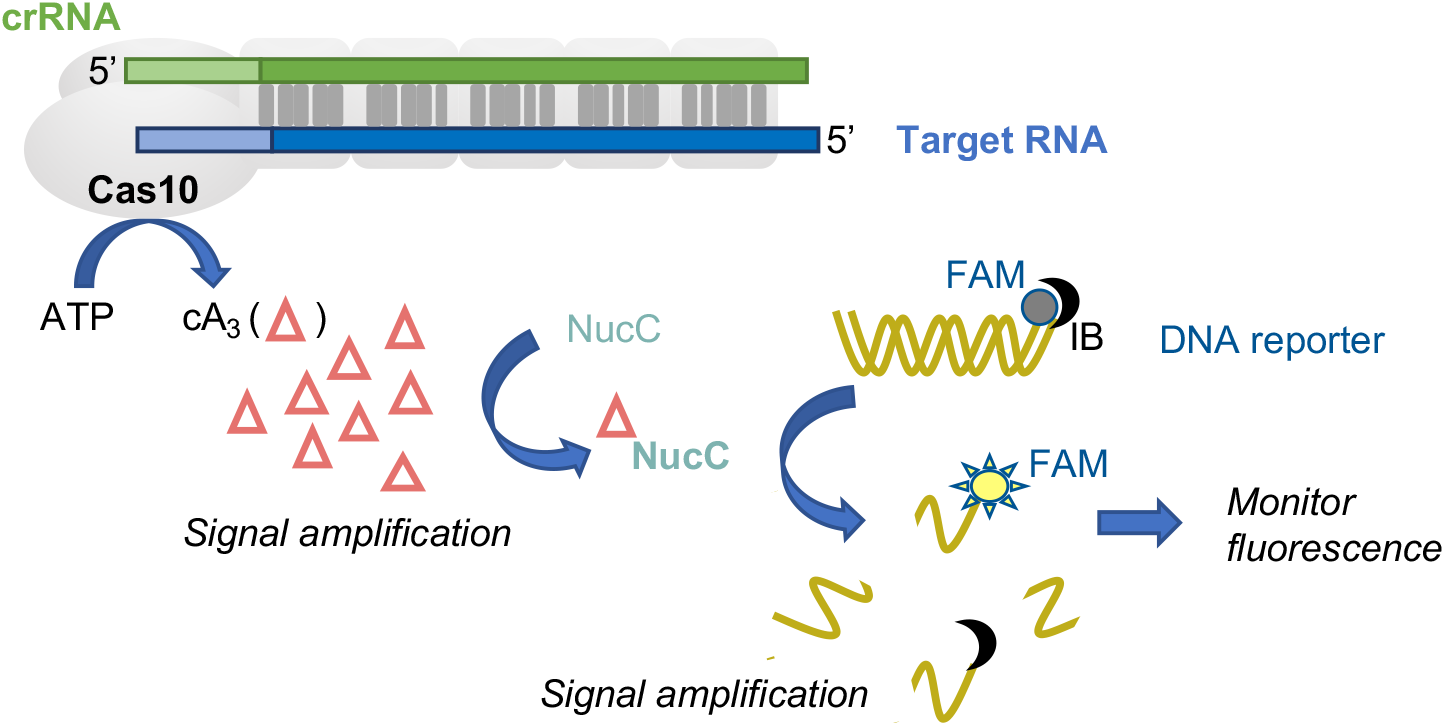
Schematic representation of the VmeCmr-NucC coupled assay. The Cas10 subunit of the VmeCmr complex produces cyclic triadenylate (cA_3_) upon binding of target RNA to the crRNA. The nuclease NucC in turn is activated by cA_3_ and starts to degrade the double-stranded reporter DNA liberating the fluorophore from its quencher. The target RNA binding event is thus coupled to a fluorescent output that can easily be monitored. FAM: 5’-6-fluorescein fluorophore; IB: 3’-Iowa Black^®^ FQ quencher.

### Specificity of target RNA detection

Type III CRISPR systems must avoid inappropriate activation by RNA targets such as anti-sense RNAs transcribed from the CRISPR locus, which could cause toxicity or even cell death. To achieve this, Type III systems sense mispairing of RNA at the Protospacer flanking site (PFS), which is immediately 3’ of the RNA duplex formed between the target RNA and the crRNA, corresponding to the repeat-derived 5’-handle of the crRNA. When an anti-sense CRISPR RNA binds, it base-pairs along the length of the PFS, preventing activation of the HD nuclease or cyclase activities of Cas10 (15–17,19,22,23,26. A detailed investigation of the Type III-B system from *Thermotoga maritima* (*Tma*Cmr) (50) revealed that the three nucleotides at positions −1 to −3 of the PFS are crucial in regulating Cas10 activity, consistent with observations in type III-A systems (51–53) and furthermore that guanine at position −1 is sensed directly, rather than via base-pairing, to keep the complex in an inactive state (50).

We first tested the importance of the PFS for VmeCmr activity (Figure 5) by changing one or more nucleotides in the target RNA sequence. At position −1, all four nucleotides were well tolerated but a guanine at position −1 resulted in a slightly higher level of activation of VmeCmr. Introduction of a U:A base pair at position −2 reduced activity significantly, but equivalent base pairs introduced singly at positions −3 to −5 had only a modest impact on activity (Figure 5C). Indeed, the presence of a U:A base pair at position −3 slightly enhanced activity. However, introduction of a run of three base-pairs at positions −2 to −4 virtually abolished VmeCmr activity, emphasising the importance of targets which do not base pair with the crRNA in this region, as observed previously (51–53).

**Figure 5.**
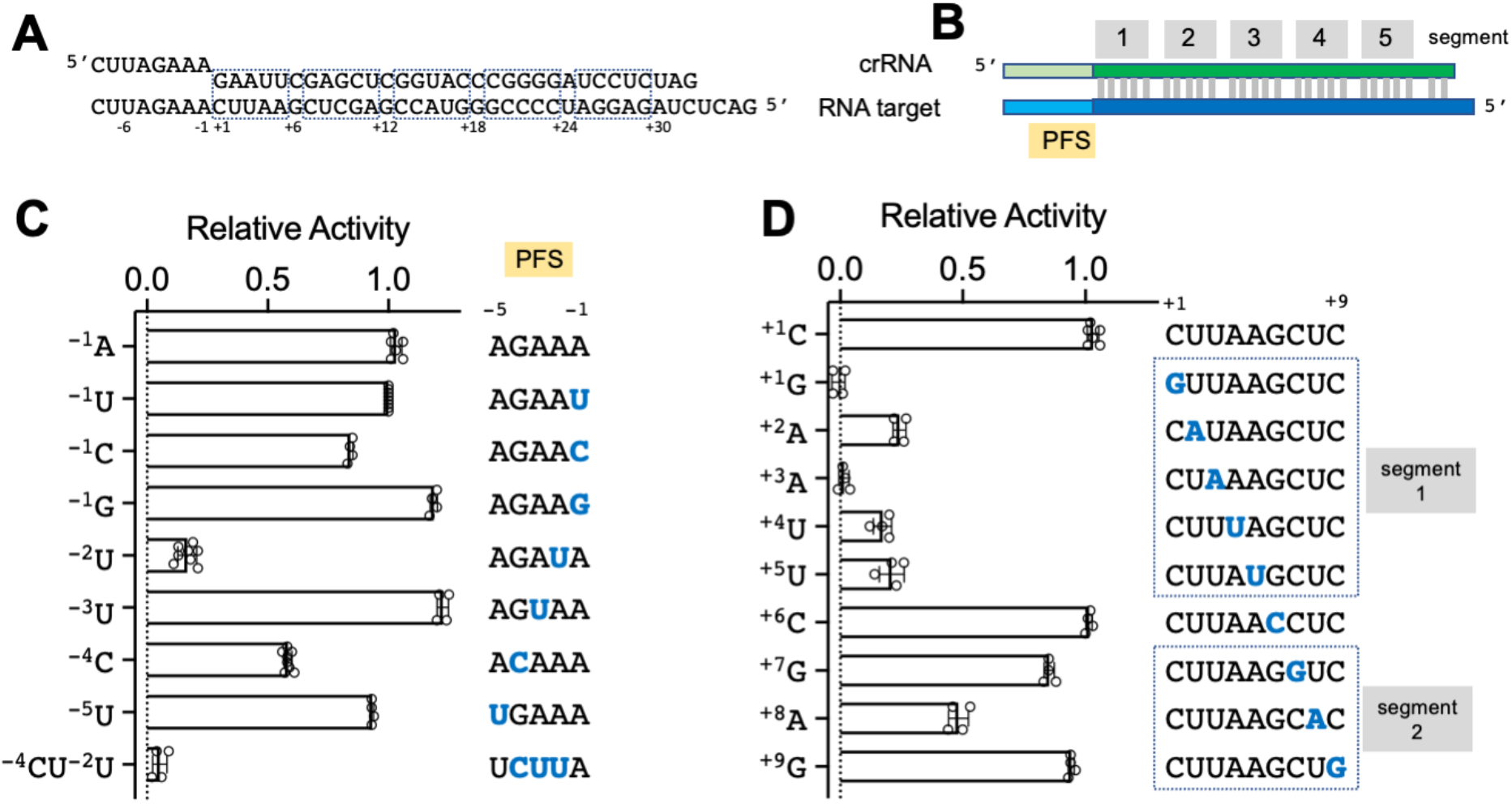
Target specificity of VmeCmr. **A:** Sequences of the crRNA (top) and synthetic target RNA (bottom). **B:** Main features of the two RNA sequences. **C:** Effect of changes in the sequence of the PFS in the target RNA on activity. **D:** Effect of mismatches in the spacer region on activity. Activity was determined relative to the reference target RNA sequence shown in (A). Nucleotides that differ from the reference sequence are shown in blue. The target RNA concentration was 5 pM.

Type III CRISPR systems are tolerant of extensive mis-pairing between crRNA and target RNA, a factor which is postulated to limit viral escape by mutation (26,50,54–57). The bound target RNA can be divided into 5 bp segments followed by a sixth nucleotide that is flipped out of the duplex by the Cas7 subunit (52,53,58,59). Segment 1, matching the 5’-end of the spacer sequence, adjacent to the PFS, is particularly important for Cas10 activation (26,50), analogous to the “seed” region next to the protospacer adjacent motif in type I, II and V systems. To investigate this, we changed single nucleotides in the target RNA at positions 1 to 9, making the nucleotide at each position identical to, rather than complementary to, the crRNA sequence (Figure 5D). At the concentration of target RNA tested (5 pM), single mutations at positions 1 to 5 all resulted in a significant decrease in cA_3_ production and therefore NucC activity. Mutation at position 6, which is not base-paired in the ribonucleoprotein complex, had no effect on the cyclase activity, as expected, and there were modest reductions in cyclase activity for positions 7 and 8 in segment 2.

VmeCmr is thus exquisitely sensitive to single nucleotide mismatches in segment 1, which contrasts with the findings for *TmaCmr*, where significant effects on Cas10 activity were only observed when four or five nucleotides were mutated simultaneously (50), but is in good agreement with findings for the *S. solfataricus* type III-D and *Staphylococcus epidermidis* type III-A complexes (26,60). A mismatch at position +1 or +3 resulted in almost complete abolition of the fluorescent signal, reflecting effective suppression of the cyclase activity of Cas10.

### Sensitive detection of the SARS-CoV-2 N gene RNA by VmeCmr/NucC

We next investigated the sensitivity of the VmeCmr/NucC system for detection of larger RNA species, using the SARS-CoV-2 N gene as an exemplar. To programme VmeCmr to detect the SARS-CoV-2 RNA specifically, we designed, expressed and purified six different VmeCmr complexes carrying guide RNAs designed to match a range of positions in the SARS-CoV-2 N gene (Figure 6A). The VmeCmr constructs were named according to the first nucleotide of the N gene matching the crRNA (Figure 6A, Table 1). Each was designed to have a G at position −1 of the PFS, as that provided the highest activity with the reference target set (Figure 5C). We generated a ~1250 nt *in vitro* transcript of the N gene to serve as target RNA for the assays.

**Figure 6.**
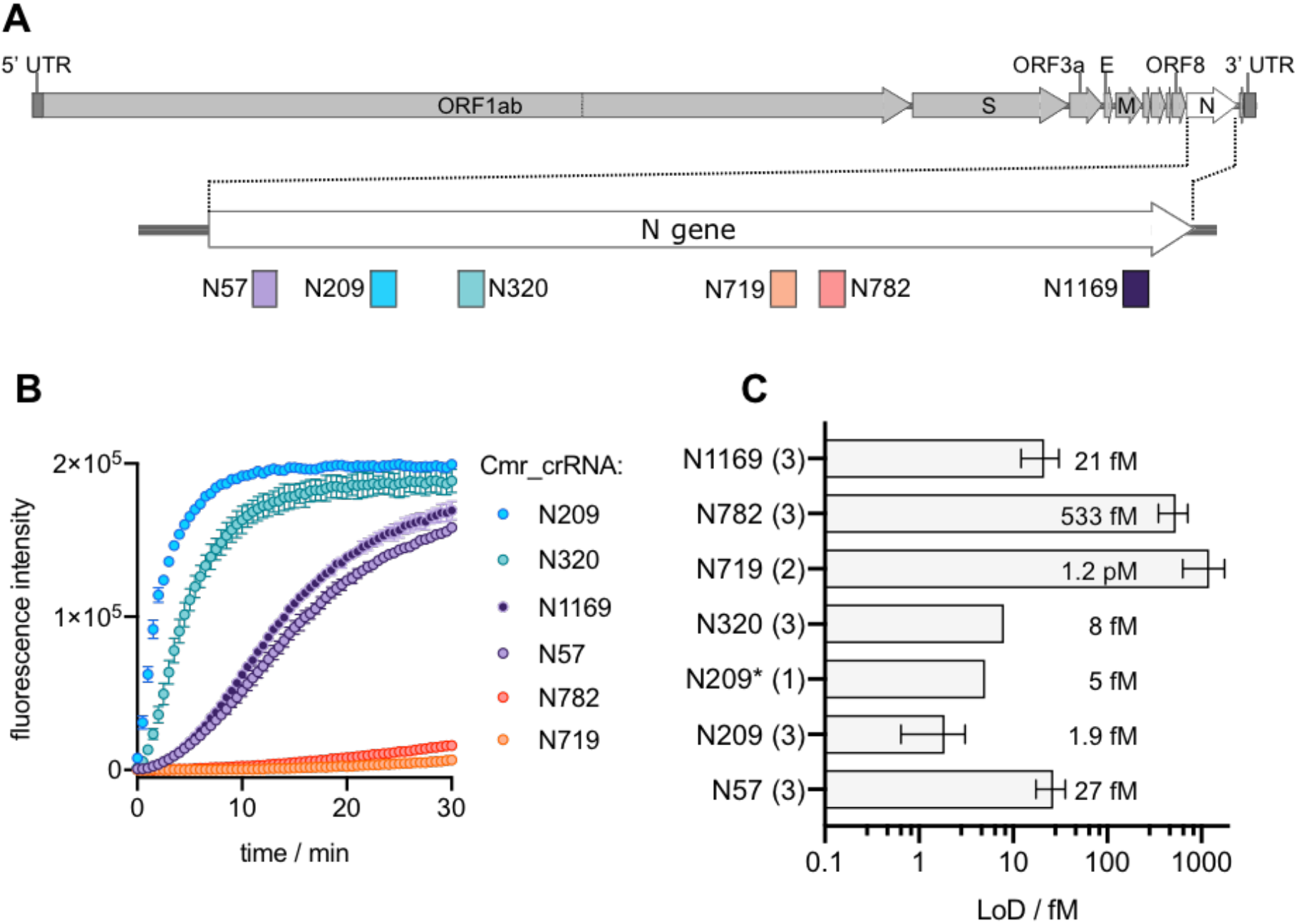
SARS-CoV-2 N gene targeting VmeCmr complexes. **A**: Representation of the SARS-CoV-2 genome and location of the VmeCmr^crRNA^ target sites on the N gene. DNA maps were created with SnapGene Viewer. **B**: VmeCmr complexes charged with crRNAs targeting different positions in the N gene responded in varying degrees to the presence of target RNA. The curves shown were obtained with 2.6 pM N gene transcript and 100 nM VmeCmr complex in the coupled NucC assay. **C**: LoDs for SARS-Cov-2 N gene-targeting VmeCmr complexes. The number of independent experiments is given in brackets next to the crRNA designation. Wild type VmeCmr complexes were used except for N209 (N209*: VmeCmr^N209^ Cmr4 D26A). LoDs were determined from the fluorescence intensities relative to a reference (no RNA) as described in Materials and Methods. Target concentrations with mean fluorescence intensities higher than the mean intensity of the reference plus 10 SDs were regarded as detected. The quoted concentrations correspond to the average LoD value. Progress curves and fluorescence intensities used to determine the LoD are provided in Figures S5 and S6, respectively.

In the course of our studies, we noted that the background activity in the absence of any added RNA target became significant for some of the six VmeCmr constructs (Figure S4A). We therefore subjected all complexes to a further purification step by heparin chromatography, which significantly reduced the background activity without affecting the signal generated by the activated complex (Figure S4B, C). Accordingly, all further assays were conducted with VmeCmr complexes that had undergone the additional heparin purification step.

As is clear from Figure 6B, we observed a wide range of activities for the six different complexes. The best ones generated a large fluorescent signal within 1-2 min when activated by 2.6 pM transcript while the least sensitive constructs gave only a marginal signal. The difference in activity was reflected in the LoD obtained for each complex (Figure 6C). The most sensitive complexes were N209 and N320 with LoDs of 1.9 fM and 8 fM, respectively. The N719 complex was the poorest with an LoD of 1.2 pM; thus, the measured LoDs spanned 3 orders of magnitude for the six investigated VmeCmr complexes. We also tested the Cmr4 D26A variant of the N209 targeting VmeCmr complex. This mutation targets the Cmr4 (Cas7) active site and is known to prevent degradation of the target RNA in all other type III systems studied (6–8,19,26,52,60. It has previously been shown that preventing target RNA degradation in type III CRISPR-Cas complexes leads to increased cOA production (21,26,60) which in turn would be expected to lower the target RNA concentration required to trigger NucC activity. For VmeCmr^N209^ Cmr4 D26A, however, no improvement in sensitivity was observed.

We reasoned that target RNA secondary structure might in part be responsible for the observed range of LoDs. To test this hypothesis, we selected three VmeCmr complexes (N719, N57 and N209, representing the highest, middle and lowest LoDs, respectively) and compared their activity when presented with either the N gene transcript or an RNA oligonucleotide containing the target sequence plus flanking region (Figure 7). The best performing N209 complex generated equivalent fluorescent signals when activated with the transcript and oligonucleotide targets at either 10 or 100 fM concentration, suggesting that this region in the N gene is equally available for VmeCmr binding in the oligonucleotide and transcript. However, the mid-performing N57 complex did exhibit a significant improvement in activation when presented with the oligonucleotide target compared to transcript. With the oligonucleotide target, the N57 complex showed a similar activity to the best performing

**Figure 7.**
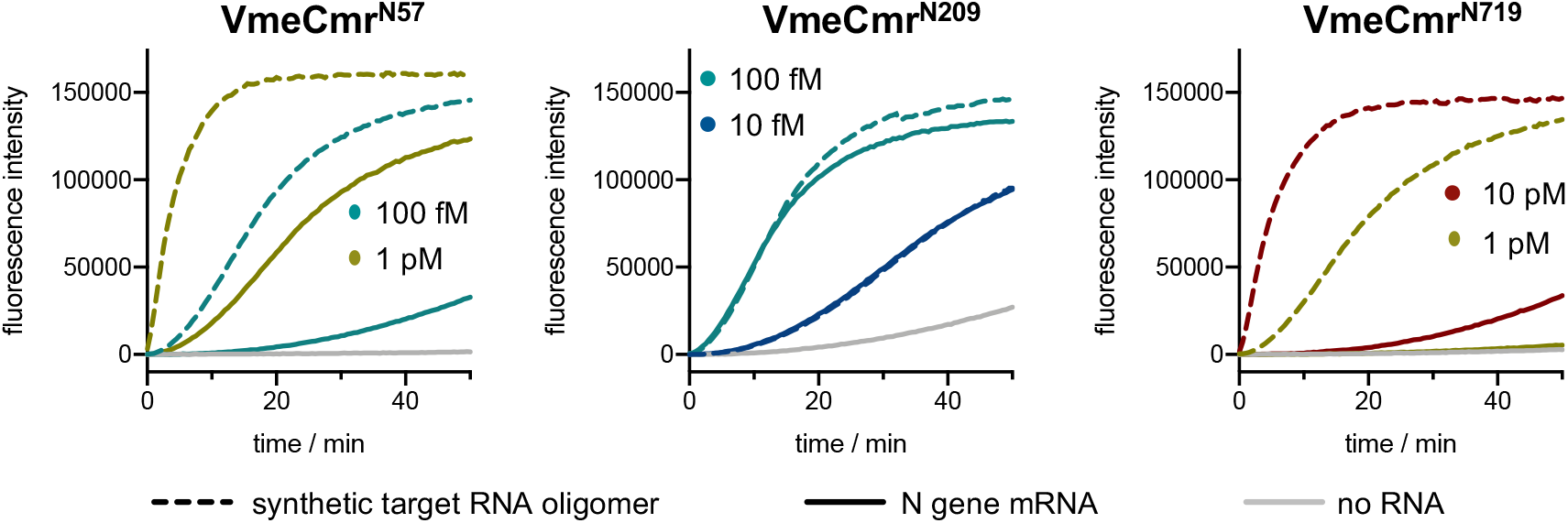
Comparison of N gene transcript and RNA oligomers as target RNA. The VmeCmr / NucC coupled assay was performed under standard conditions with 50 nM N209, 100 nM N57 or 100 nM N719 and either N gene transcript (solid line) or the 44 nt synthetic RNA (dashed line) containing the cognate target at the indicated concentrations. Identical target concentrations have the same colour.

N209 complex (Figure 7). This is consistent with the possibility that the N57 target site in the RNA transcript is partially obscured by the secondary structure of the RNA. Finally, the N719 complex was much more active against the oligonucleotide target than the transcript, but still did not approach the sensitivity of the other two complexes studied, suggesting that RNA secondary structure is not the only variable that influences VmeCmr activity.

Finally, we tested our best N gene-targeting VmeCmr complex, N209, against SARS-CoV-2 viral RNA (Figure 8, Figure S7). SARS-CoV-2 was propagated by infecting Vero E6 cells and culture supernatant was harvested to give viral stock at 6·10^6^ PFU ml^−1^. The viral stock was 10-fold serially diluted and polyA RNA was added as carrier RNA to each sample before RNA extraction using a commercial kit to give extracts 1 (highest concentration) to 6 (lowest concentration). The extracts were diluted 20-fold in triplicate into our coupled VmeCmr^N209^ / NucC assay. Both the wild type and the Cmr4 D26A (cas7 mutant) complexes were activated by SARS-CoV-2 RNA extracted from 6·10^3^ PFU ml^−1^ – 6·10^6^ PFU ml^−1^ stocks (Extracts 4 – 1) in the presence of polyA carrier RNA. As it is likely that many more RNA molecules than viable viral particles existed in the viral stock, we sought to estimate the N gene concentration in the extracts by comparison to a standard curve of known N gene transcript concentrations using RT-qPCR (Figure S8, Table S2). In this manner, we obtained a concentration of 8 fM for extract 4, the lowest dilution to yield a clear signal in the assay. Extract 5, at < 1 fM concentration, was not reliably detected. This is consistent with the LoD for the N gene transcript of 2 fM suggesting that VmeCmr can detect target RNA with high sensitivity in complex mixtures of nucleic acids. There was no significant difference between the wild type and Cmr4 D26A variant complexes.

**Figure 8.**
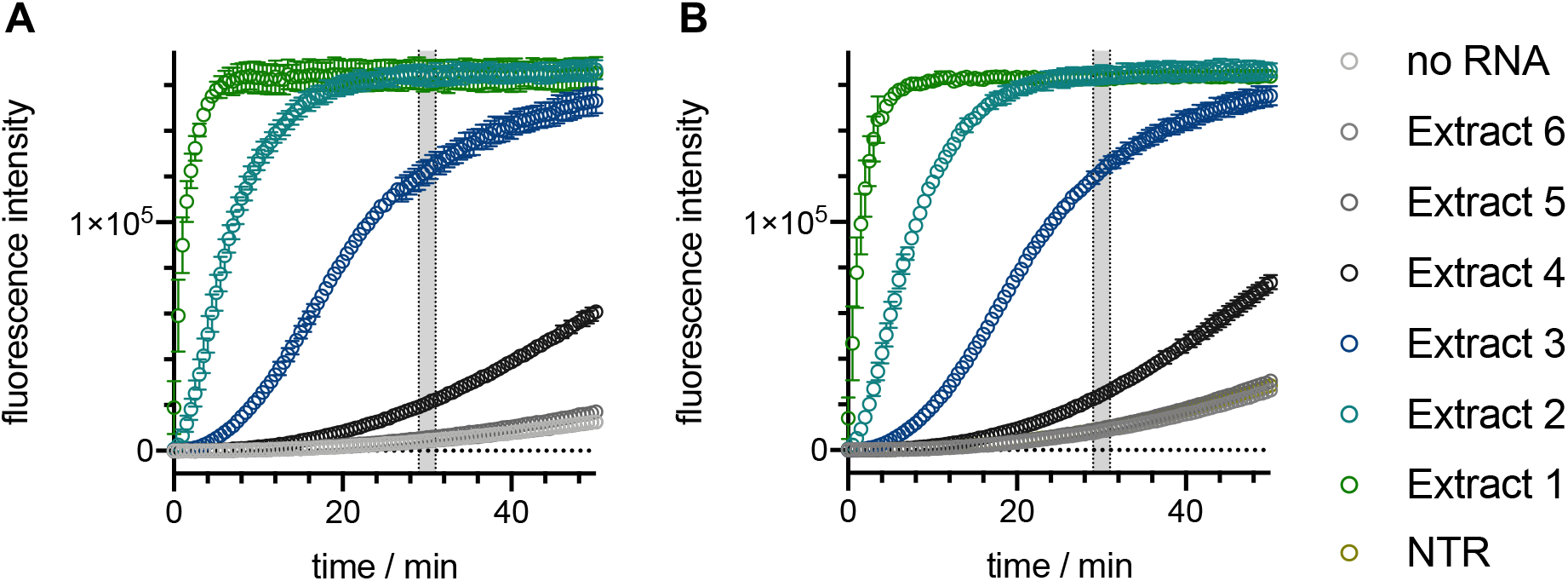
Detection of SARS-CoV-2 N gene in viral extracts. Extracts 1 – 6 were obtained by extracting a 10-fold serial dilution of viral stocks ranging from 6·10^6^ to 6·10^1^ PFU ml^−1^, respectively. The assay was performed under standard conditions in triplicate. **A**: 50 nM wild type VmeCmr^N209^; **B**: 25 nM VmeCmr^N209^ Cmr4 D26A. NTR: non-target RNA.

## Discussion

Here, we investigated a type III-B CRISPR system from *V. metoecus*. The locus is highly conserved in strains of *V. cholerae* and *V. metoecus* sampled over many years and is always prophage-encoded (34). It has been described as a hybrid III-B/I-F system as it a type III-B effector with a type I-F Cas6 and CRISPR sequence (34). The associated CRISPR locus varies in length and content, and some examples have spacers with 100% identical matches to *Vibrio* plasmids, prophage and phage genes, suggesting that this CRISPR system may play a role in inter-phage conflict (62,63). These spacers share a CRISPR type I-F 3’-GG PAM, which is not characteristic of “pure” type III systems (34). It remains to be determined whether this system functions with a CRISPR-type immunity mechanism, clearing invading phage, or with a CBASS/NucC-like abortive infection mechanism (35), killing the infected cell.

This CRISPR system provides immunity by detecting target RNA and generating a cA_3_ signalling molecule which activates the associated NucC endonuclease. The LoD is observed with 10 pM synthetic cA_3_ for NucC alone, and close to 1 fM target RNA when coupled to the most sensitive VmeCmr complex, suggesting there is at least a 1000-fold signal amplification due to cA_3_ synthesis, which is in line with previous studies of other systems (46). Analysis of the sequence specificity of VmeCmr for target RNA revealed a high degree of discrimination of single nucleotide mismatches between the crRNA and target RNA in segment 1, demonstrating that VmeCmr can function as a highly sequence-specific defence system. The tolerance observed for mismatches outwith segment 1 reflects a plasticity that is shared by other type III systems (26,50,54–57), reducing the ability of viruses to mutate target sites to avoid detection by CRISPR defence (54). In the protospacer flanking sequence (PFS), extensive base-pairing at positions −2 to −5 limits activation of Cas10, a common property for type III systems that ensures anti-sense RNA generated from the CRISPR locus does not activate type III CRISPR defence. We observed a modest sequence preference at the −1 position where a G is slightly favoured over other nucleotides whilst other systems do not discriminate at the −1 position (15,54) and one, TmaCmr, is strongly inhibited by a G at position −1 (50). Notably, this preference for a G at position −1 fits with the presence of a 3’-GG PAM in the protospacers present in the CRISPR locus (34). For VmeCmr, a single base pair at position −2 in the PFS is sufficient to severely reduce Cas10 activation whilst other systems tend to tolerate single base pairs in positions −2 through −5 (50,53,54).

Taking into account the lessons learned from the reference RNA target, we designed six guide RNAs for the detection of the SARS-CoV-2 N gene transcript. This revealed stark differences in the sensitivity, ranging over 3 orders of magnitude, between the best and worst performing guide RNAs. By comparing the sensitivity of the system when presented with short RNA oligos versus larger transcripts, we confirmed that RNA secondary structure is an important factor for VmeCmr, with the worst performing complex N719 improved by two orders of magnitude when provided with an RNA oligonucleotide target rather than a transcript (Figure 7). This is a familiar concept in molecular biology, where the availability of RNA target sequences can be modulated by changes in RNA secondary structure – for example in riboswitches (reviewed in (64)). Indeed, wide variations in the sensitivity of the Cas13 effector have been observed with different guide RNA sequences during SARS-CoV-2 assay development (38). In contrast, a recent study from the Wiedenheft lab (40) detected only minor differences in the sensitivity of 10 guides designed to detect different regions of the N gene using a type III effector. A notable difference is that our assays were carried out at 37 °C whereas the Wiedenheft group used a thermostable TtCsm complex active at 60 °C. It is perhaps not entirely coincidental that type III CRISPR systems predominate in thermophiles (65), where the secondary structure of RNA targets might not present so much of a problem.

One unexpected finding was that the background cyclase activity of VmeCmr in the absence of any target RNA varies between constructs and is proportional to the maximal activity when target RNA is present (Figure S4A). Thus, the complex with the N209 guide had a much higher background activity than the one with N719. This background rate was strongly reduced by heparin chromatography and a concomitant decrease in the A260/A280 ratio of the purified complexes. One possibility is that *E. coli* mRNA with partial matches to the crRNA sequence can co-purify with the VmeCmr complex and are sufficient to provide partial activation of the cyclase.

The combination of VmeCmr and NucC with an optimal guide RNA and fluorescence readout provided a LoD of 2 fM for detection of the SARS-CoV-2 N gene transcript, consistent with the observation that a reaction with fewer than 1 × 10^5^ copies of the viral RNA (8 fM) can be readily detected (Figure 8). This compares with a LoD using the *T. thermophilus* Csm (TtCsm) system of 1 × 10^7^ copies (100 pM) (40) and 1.2 × 10^10^ copies (1 nM) detected by the TtCmr system (41). The >100-fold higher sensitivity of the assay we describe here is likely due to a combination of factors. These include the high affinity of NucC for cA_3_, low background activity of the nuclease, low levels of ring nuclease activity, high levels of cA_3_ production by VmeCmr and the use of a dsDNA species as a molecular beacon rather than a less stable RNA reporter and screening for optimal guide RNAs. We observed little difference between the wild-type VmeCmr complex and the Cmr4 D26A variant in our assays, despite the fact that this mutation abolishes the ability to degrade bound target RNA (7). One potential explanation is that we are approaching the limits of target RNA binding affinity instrinsic to the Cmr complex, but this remains an open question for future study.

Our work provides a proof of concept for the use of cA_3_ generating type III CRISPR systems coupled to the NucC effector nuclease as the basis for a rapid, sensitive and specific assay for any desired RNA sequence. The sensitivity of our best performing complex is on a par with first generation Cas13 assays; by way of comparison, Lbu-Cas13a programmed with a single guide RNA has a LoD of 10 fM of target RNA (38,66). There is clear potential for further improvement, for example by optimising and multiplexing guide RNAs, which has yielded clear benefits with Cas13a (38). Approaches that reduce the secondary structure of RNA samples are also likely to provide benefits in sensitivity. The signal generated by NucC could also be readily adapted to alternative assay formats such as lateral flow. Complexities arising from sample complexity (for example detection of RNA in saliva samples) have yet to be explored but would be expected to reduce assay sensitivity – there remains the option to couple the VmeCmr/NucC assay to an isothermal amplification step to boost the signal to noise ratio.

## Acknowledgements

Thanks to Dr Januka Athukoralage and Dr Jane Hilton for scientific discussions and technical assistance.

## Author contributions

S.G. designed experiments, carried out experiments and analysed data in consultation with M.F.W. and C.S.A. C.S.A. cultured and purified the SARS-CoV-2 virus and purified the RNA. M.F.W. conceptualised the project, obtained funding, carried out formal analysis and prepared the original draft of the manuscript. All authors contributed to writing, review and editing.

