## Supplementary data for "Specificity and sensitivity of an RNA targeting type III CRISPR complex coupled with a NucC endonuclease effector"

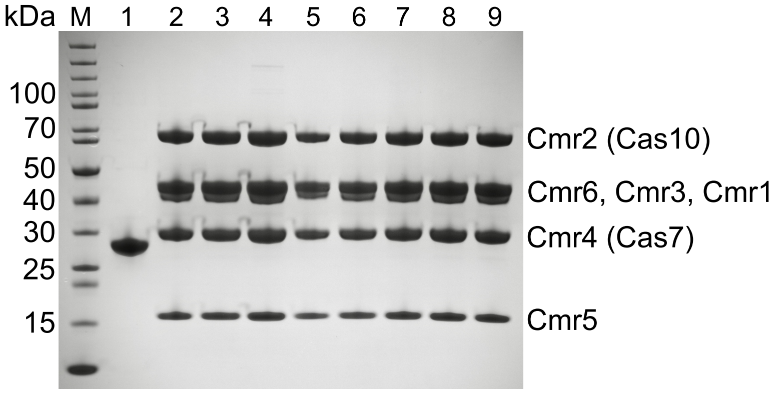


**Figure S1.SDS-PAGE of purified VmeCmr complexes and VmeNucC used in assays.** M: PageRuler Unstained (Thermo Scientific) protein marker; 1: VmeNucC; 2: VmeCmr$^{pUC}$; 3: VmeCmr$^{N57}$; 4: VmeCmr$^{N209}$; 5: VmeCmr$^{N209}$ Cmr4 D26A; 6: VmeCmr$^{N320}$; 7: VmeCmr$^{N719}$; 8: VmeCmr$^{N782}$; 9: VmeCmr$^{N1169}$. The crRNA target for each Cmr complex is indicated as superscript.

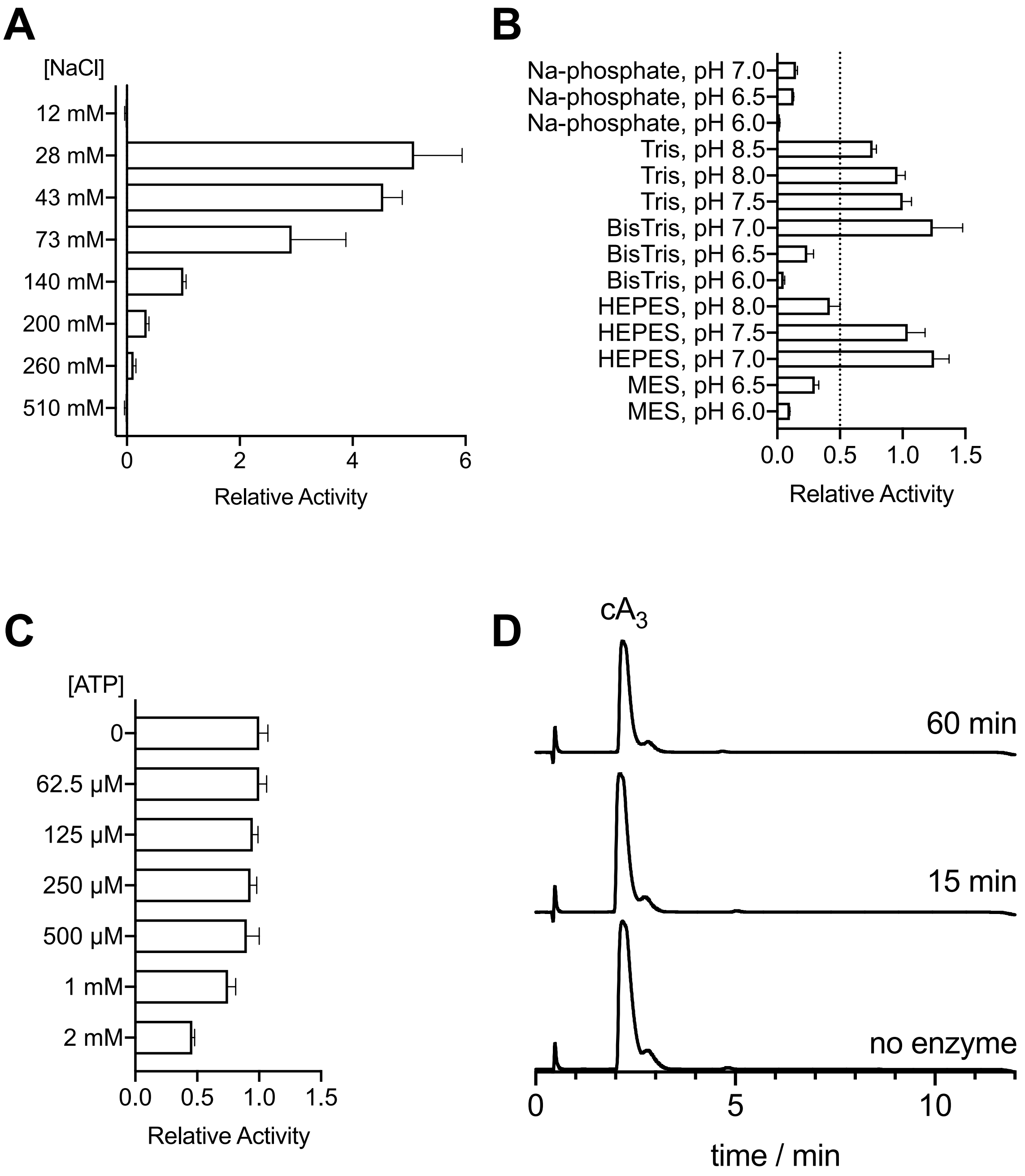


**Figure S2.Factors influencing NucC activity.** Relative activities were determined by comparing the rates of the reactions relative to an arbitrarily chosen reference condition. Rates were assumed to be proportional to the change in fluorescence signal over time and calculated from the slopes of linear regression analysis. A: Effect of NaCl concentration. B: NucC activity in different buffers. C: Effect of ATP on NucC nuclease activity. D: LC chromatograms with UV monitoring at 254 nm showing the reaction of 100 µM $cA_3$ with 0.5 µM NucC under assay conditions after 15 and 60 min. A reaction in which NucC had been omitted was allowed to proceed for 60 min.

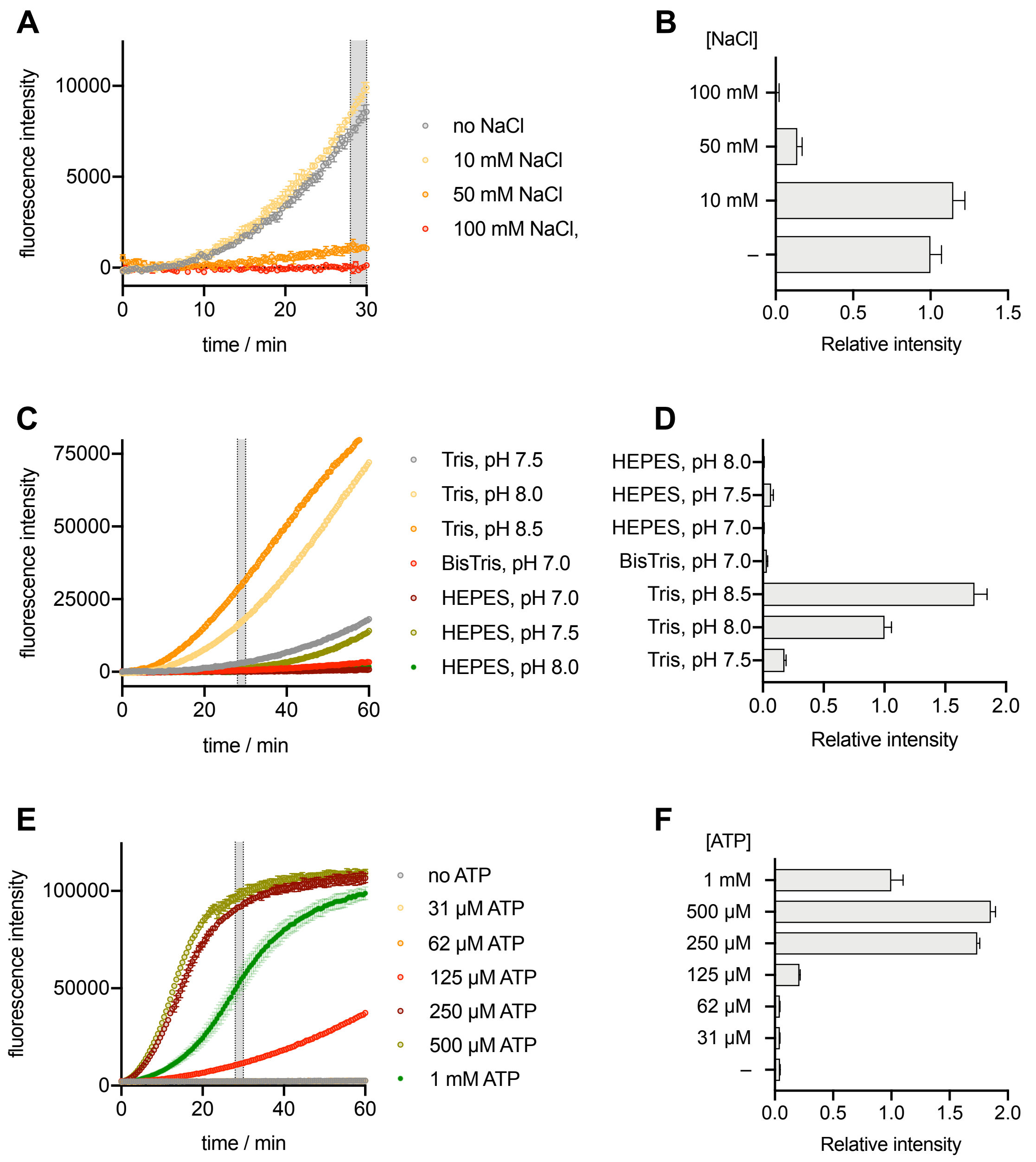


**Figure S3. Screening of reaction conditions for the coupled VmeCmr/NucC assay.** A, C, E: Progress curves. B, D, F: Fluorescence intensities extracted from corresponding progress curves (28 – 30 min, shaded in grey) relative to an arbitrarily chosen reference condition. A and C were baseline corrected using the progress curves obtained in the absence of target; no baseline correction was performed for E. Reaction conditions for A and B: 100 nM Cmr$^{pUC}$, 200 pM pUC target, 50 mM Tris-HCl (pH 7.5), 6 % (v/v) glycerol, 10 mM $MgCl_2$, 250 µM ATP, 12 mM NaCl. Additional NaCl was supplemented as indicated in the graphs; C and D: 100 nM Cmr$^{pUC}$, 200 pM pUC target, 50 mM buffer as indicated, 22 mM NaCl, 6 % glycerol, 10 mM $MgCl_2$, 250 µM ATP; E and F: 100 nM Cmr$^{pUC}$, 200 pM pUC target, 12.5 mM HEPES (pH 7.5), 12 mM NaCl, 6 % glycerol, 10 mM $MgCl_2$, ATP as indicated. All reactions contained 250 nM NucC and 125 nM NucC substrate. They were preincubated for 10 min at 37 °C before addition of NucC. Assays were performed in duplicate

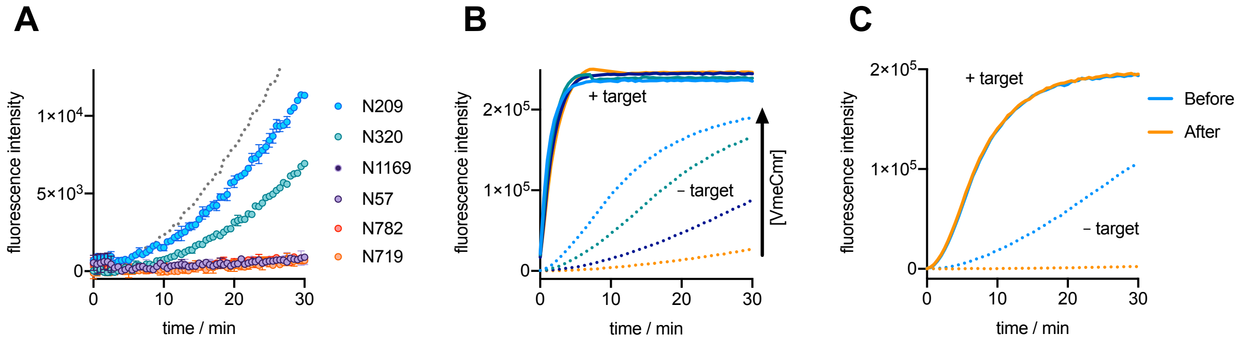


**Figure S4. Factors influencing unregulated $cA_3$ synthesis in VmeCmr complexes.** Progress curves of the coupled VmeCmr/NucC assay are shown. Only the mean of duplicate reactions is shown for clarity (B, C). N gene transcript was used as target RNA. **A**: VmeCmr / NucC background activity in the absence of target RNA. Companion graph to Figure 6B. The dotted curve corresponds to N782 in the presence of 2.6 pM target RNA (provided for scale). **B**: Reducing the VmeCmr concentration can reduce background activity (in the absence of target RNA) without affecting target-induced $cA_3$ synthesis. $VmrCmr^{N209}$ before heparin purification was used at 31 – 250 nM. **C**: Further purification by heparin chromatography significantly reduced the background activity without affecting target-induced activity (100 nM $VmeCmr^{N320}$).

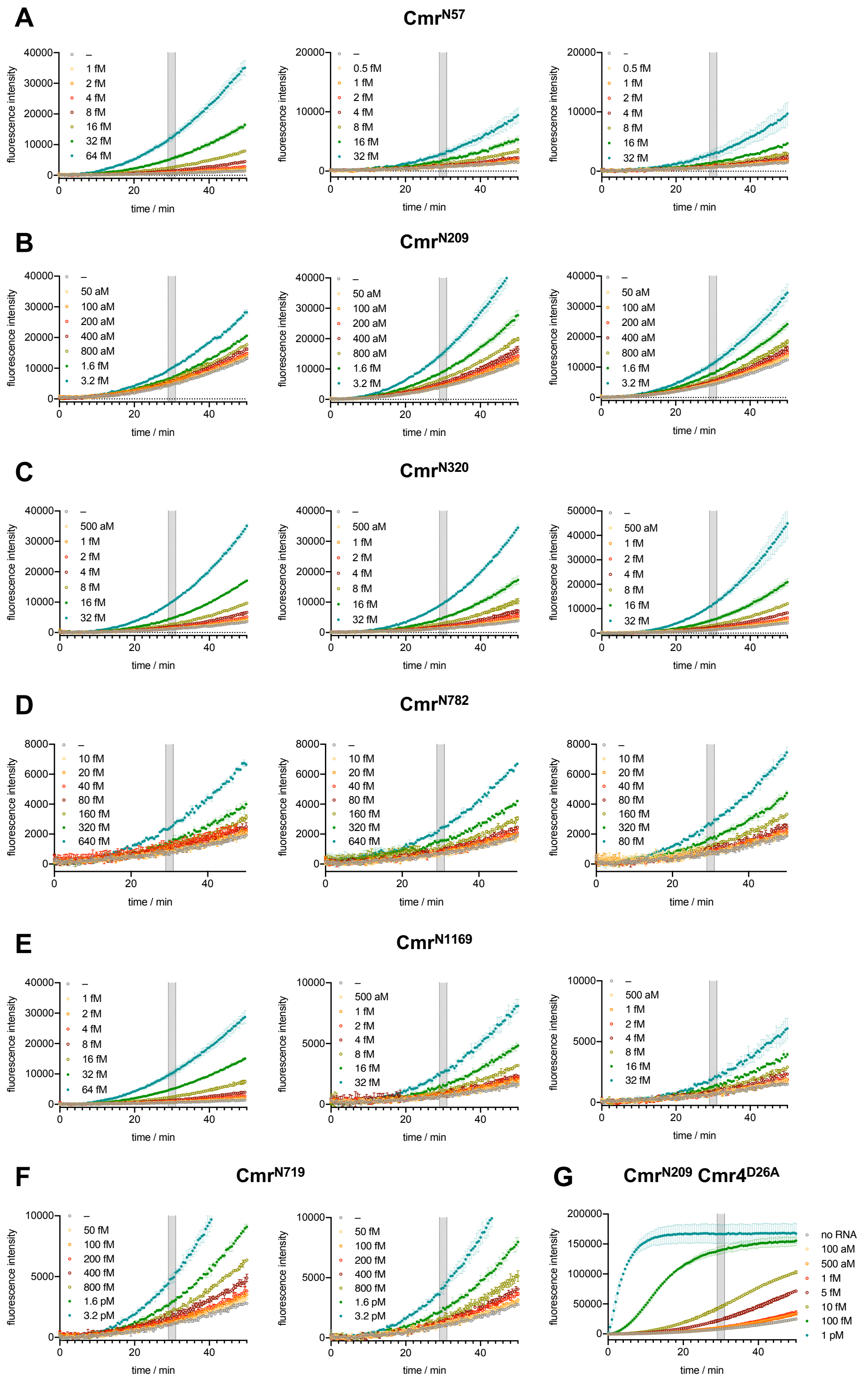


**Figure S5. Progress curves used to determine the LoD of SARS-Cov-2-targeting VmeCmr complexes.** The mean and standard deviation of the fluorescence intensities are shown. Standard assay conditions as described in Materials and Methods were used. The Cmr concentrations were 100 nM for A (Cmr$^{N57}$), D (Cmr$^{N782}$), E (Cmr$^{N1169}$), and F (Cmr$^{N719}$), 50 nM for B (Cmr$^{N209}$), and C (Cmr$^{N320}$), 25 nM for G (Cmr$^{N209}$ Cmr4 D26A). N gene transcript at indicated concentration ranges was used as target throughout. The fluorescence intensities between 29 – 31 min (shaded in grey, Figure S6) were used to determine the LoD. Each independent experiment was performed in duplicate.

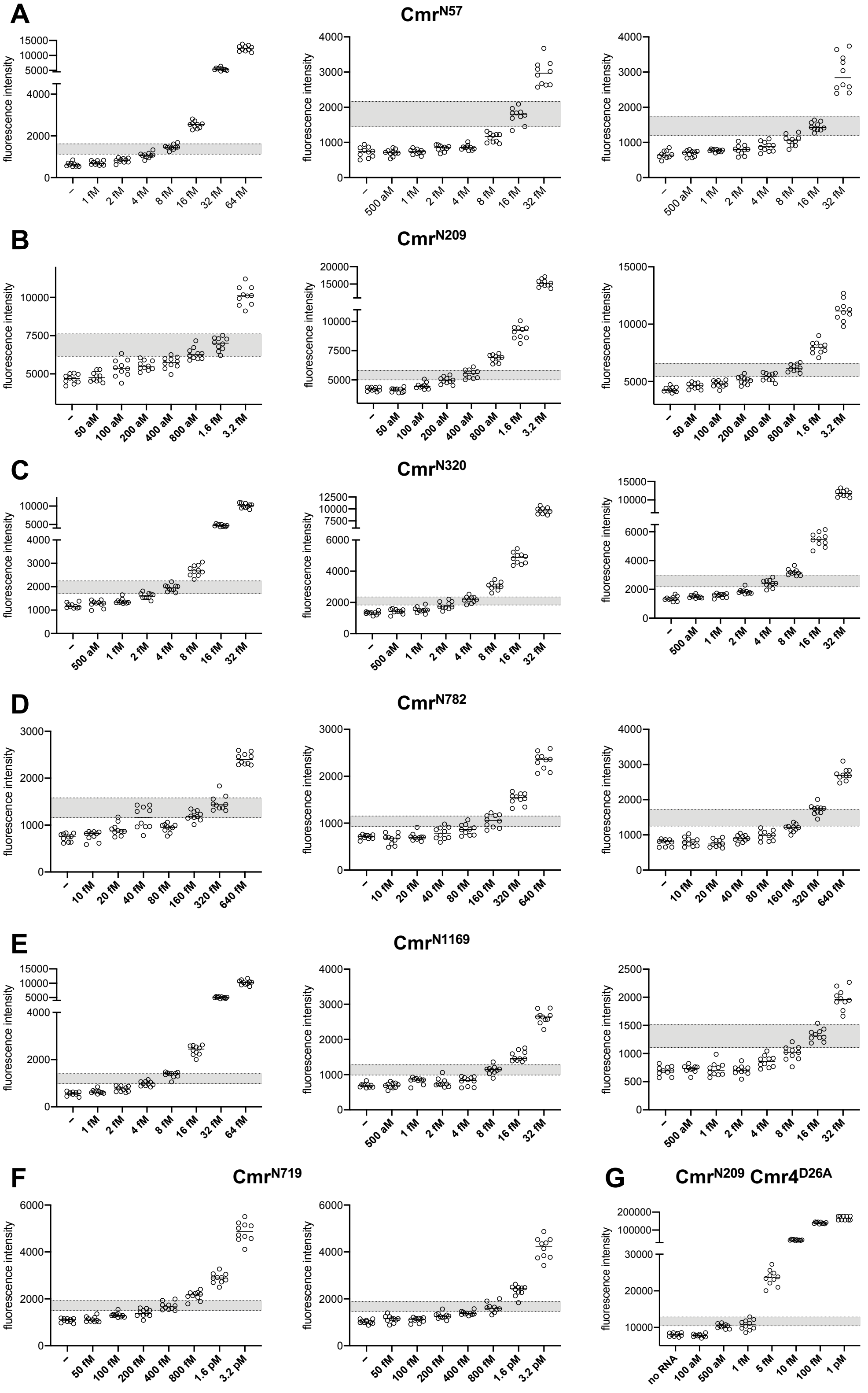


**Figure S6. Fluorescence intensities used to determine the LoD of SARS-CoV-2-targeting VmeCmr complexes.** The fluorescence intensities between 29 – 31 min are shown as individual values and mean (**Error! Reference source not found.**). The shaded area corresponds to *y* values between 5 and 10 standard deviations greater than the mean of the reference (no RNA). The lowest measured target concentration (mean) above the shaded area was taken as the LoD. For clarity, the *y* axis has been split for some experiments and the scaling of the two sections is not identical.

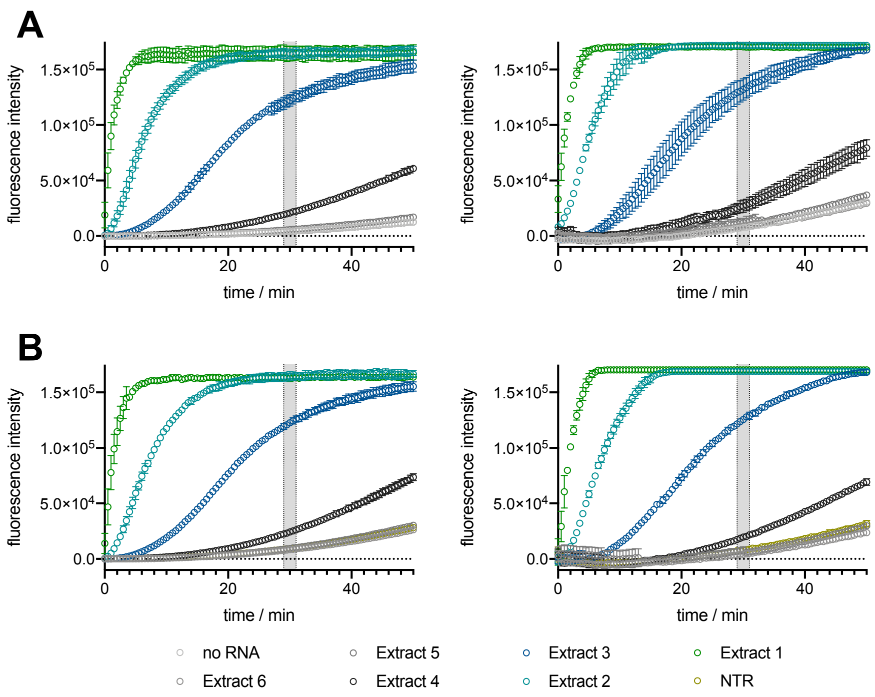


**Figure S7. Detection of SARS-CoV-2 N gene in viral extracts.** Extracts 1 – 6 were obtained by extracting a 10-fold serial dilution of viral stocks ranging from $6 \cdot 10^6$ to $6 \cdot 10^1$ PFU $ml^{-1}$, respectively. The assay was performed under standard conditions in triplicate. **A**: 50 nM wild type $VmeCmr^{N209}$; **B**: 25 nM $VmeCmr^{N209}$ Cmr4 D26A. NTR: non-target RNA. Complete data set for Figure 8. Two different batches of SARS-CoV-2 RNA were used in each set.

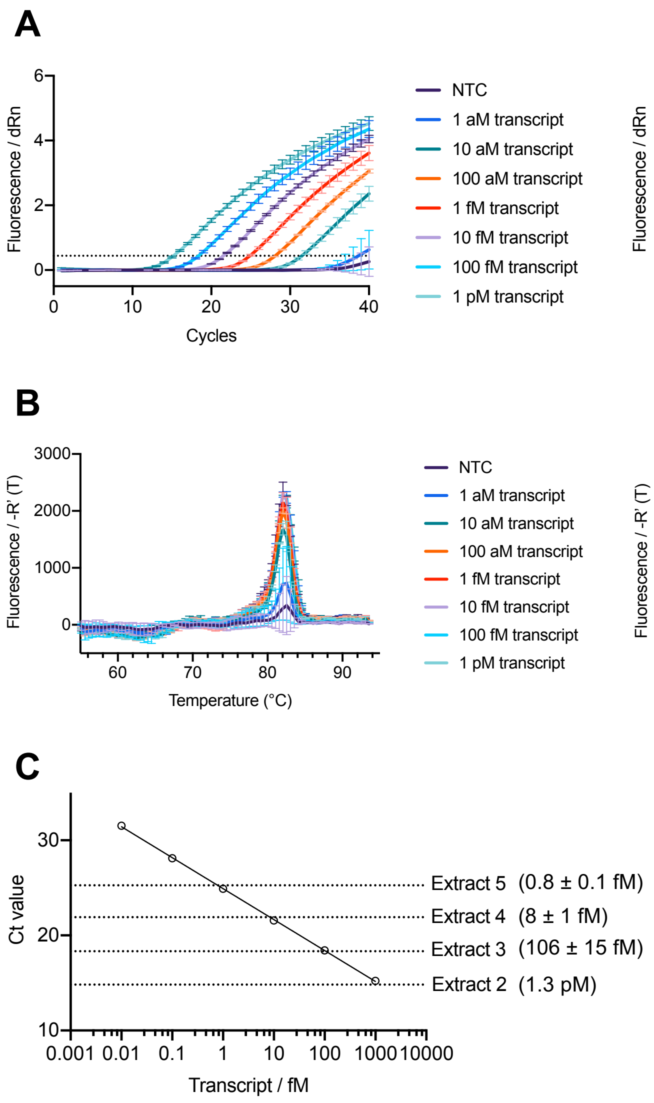


**Figure S8. RT-qPCR analysis of SARS-CoV-2 N gene transcript and genomic RNA extracts. A**: Amplification plots, the threshold value is indicated by a dotted line; **B**: dissociation curves; **C**: standard curve based on transcript dilution series (10 aM – 1 pM, $R^2$ 0.9999), the Ct values of the extracts are indicated by dotted lines; the concentrations were calculated from non-linear regression (semi-log line, Prism). NTC: no template control; NRT: no reverse transcriptase control. No DNA contamination was detected in the tested SARS-CoV-2 genomic RNA extracts. Only the highest N gene transcript concentration was tested for DNA contamination; even though detectable levels of DNA were present in this sample it was deemed negligible ($\Delta$Ct = -18.4 corresponding to $10^5$ – $10^6$-fold excess of RNA over DNA).

### Table S1. Data sets used for determination of LoDs.

| | | | | | | | | |
|---|---|---|---|---|---|---|---|---|
| **pUC (1)** | no RNA | NTR | 78.1 fM | 156.3 fM | 312.5 fM | 625 fM | 1.25 pM | 2.5 pM |
| mean (N = 10) | 15073 | 15599 | 20414 | 22805 | 30846 | 48161 | 72971 | 92436 |
| SD | 832 | 855 | 1145 | 1266 | 1483 | 1787 | 1668 | 1254 |
| Threshold | 23395 | 24151 | | | | | | |
| Sample above threshold? | Ref | | nd | nd | detected | detected | detected | detected |
| | | Ref | nd | nd | detected | detected | detected | detected |
| **pUC (1)** | no RNA | NTR | 78.1 fM | 156.3 fM | 312.5 fM | 625 fM | 1.25 pM | 2.5 pM |
| mean (N = 10) | 13455 | 14689 | 15338 | 15568 | 21071 | 31320 | 47989 | 66849 |
| SD | 1078 | 988 | 823 | 924 | 1213 | 2015 | 2830 | 3270 |
| Threshold | 24233 | 24569 | | | | | | |
| Sample above threshold? | Ref | | nd | nd | nd | detected | detected | detected |
| | | Ref | nd | nd | nd | detected | detected | detected |
| **N57 (1)** | no RNA | 1 fM | 2 fM | 4 fM | 8 fM | 16 fM | 32 fM | 64 fM |
| mean (N = 10) | 629 | 705 | 826 | 1070 | 1467 | 2529 | 5434 | 12340 |
| SD | 99 | 95 | 106 | 134 | 123 | 164 | 459 | 952 |
| Threshold | 1617 | | | | | | | |
| Sample above threshold? | | nd | nd | nd | nd | detected | detected | detected |
| **N57 (2)** | no RNA | 500 aM | 1 fM | 2 fM | 4 fM | 8 fM | 16 fM | 32 fM |
| mean (N = 10) | 724 | 708 | 735 | 839 | 856 | 1135 | 1757 | 2965 |
| SD | 144 | 95 | 75 | 85 | 73 | 128 | 224 | 352 |
| Threshold | 2162 | | | | | | | |
| Sample above threshold? | | nd | nd | nd | nd | nd | nd | detected |
| **N57 (3)** | no RNA | 500 aM | 1 fM | 2 fM | 4 fM | 8 fM | 16 fM | 32 fM |
| mean (N = 10) | 656 | 686 | 770 | 789 | 883 | 1058 | 1452 | 2968 |
| SD | 109 | 86 | 35 | 139 | 140 | 152 | 120 | 517 |
| Threshold | 1746 | | | | | | | |
| Sample above threshold? | | nd | nd | nd | nd | nd | nd | detected |
| **N209 (1)** | no RNA | 50 aM | 100 aM | 200 aM | 400 aM | 800 aM | 1.6 fM | 3.2 fM |
| mean (N = 10) | 4680 | 4802 | 5373 | 5543 | 5723 | 6321 | 6958 | 10096 |
| SD | 293 | 315 | 568 | 315 | 387 | 392 | 428 | 623 |
| Threshold | 7613 | | | | | | | |
| Sample above threshold? | | nd | nd | nd | nd | nd | nd | detected |
| **N209 (2)** | no RNA | 50 aM | 100 aM | 200 aM | 400 aM | 800 aM | 1.6 fM | 3.2 fM |
| mean (N = 10) | 4214 | 4137 | 4499 | 4961 | 5522 | 6839 | 9089 | 15256 |
| SD | 158 | 178 | 273 | 296 | 375 | 319 | 575 | 1091 |
| Threshold | 5793 | | | | | | | |
| Sample above threshold? | | nd | nd | nd | nd | detected | detected | detected |
| **N209 (3)** | no RNA | 50 aM | 100 aM | 200 aM | 400 aM | 800 aM | 1.6 fM | 3.2 fM |
| mean (N = 10) | 4305 | 4617 | 4798 | 5116 | 5405 | 6180 | 7968 | 11176 |
| SD | 225 | 254 | 284 | 344 | 336 | 322 | 540 | 886 |
| Threshold | 6557 | | | | | | | |
| Sample above threshold? | | nd | nd | nd | nd | nd | detected | detected |
| **N209 $Cmr4^{D26A}$** | no RNA | 100 aM | 500 aM | 1 fM | 5 fM | 10 fM | 100 fM | 1 pM |
| mean (N = 10) | 8021 | 7793 | 10357 | 10717 | 23527 | 44994 | 139748 | 166704 |
| SD | 482 | 450 | 608 | 1378 | 2134 | 2922 | 4861 | 11507 |
| Threshold | 12845 | | | | | | | |
| Sample above threshold? | | nd | nd | nd | detected | detected | detected | detected |
| **N320 (1)** | no RNA | 500 aM | 1 fM | 2 fM | 4 fM | 8 fM | 16 fM | 32 fM |
| mean (N = 10) | 1187 | 1267 | 1376 | 1611 | 1946 | 2693 | 4733 | 10115 |
| SD | 106 | 149 | 110 | 125 | 140 | 226 | 297 | 650 |
| Threshold | 2249 | | | | | | | |
| Sample above threshold? | | nd | nd | nd | nd | detected | detected | detected |
| **N320 (2)** | no RNA | 500 aM | 1 fM | 2 fM | 4 fM | 8 fM | 16 fM | 32 fM |
| mean (N = 10) | 1320 | 1426 | 1525 | 1810 | 2179 | 3053 | 4840 | 9625 |
| SD | 103 | 150 | 183 | 244 | 204 | 252 | 358 | 642 |
| Threshold | 2346 | | | | | | | |
| Sample above threshold? | | nd | nd | nd | nd | detected | detected | detected |
| **N320 (3)** | no RNA | 500 aM | 1 fM | 2 fM | 4 fM | 8 fM | 16 fM | 32 fM |
| mean (N = 10) | 1358 | 1497 | 1572 | 1851 | 2401 | 3186 | 5460 | 11788 |
| SD | 163 | 106 | 134 | 174 | 277 | 222 | 467 | 873 |
| Threshold | 2990 | | | | | | | |
| Sample above threshold? | | nd | nd | nd | nd | detected | detected | detected |

**Table S1** cont.

| | | | | | | | | |
|---|---|---|---|---|---|---|---|---|
| **N719 (1)** | no RNA | 10 fM | 20 fM | 40 fM | 80 fM | 160 fM | 320 fM | 640 fM |
| mean (N = 10) | 990 | 1022 | 1160 | 1092 | 1162 | 1151 | 1288 | 1507 |
| SD | 106 | 114 | 97 | 113 | 132 | 153 | 139 | 114 |
| Threshold | 2050 | | | | | | | |
| Sample above threshold? | | nd | nd | nd | nd | nd | nd | nd |
| **N719 (2)** | no RNA | 50 fM | 100 fM | 200 fM | 400 fM | 800 fM | 1.6 pM | 3.2 pM |
| mean (N = 10) | 1088 | 1128 | 1306 | 1394 | 1690 | 2135 | 2881 | 4862 |
| SD | 83 | 113 | 96 | 163 | 168 | 185 | 219 | 413 |
| Threshold | 1923 | | | | | | | |
| Sample above threshold? | | nd | nd | nd | nd | detected | detected | detected |
| **N719 (3)** | no RNA | 50 fM | 100 fM | 200 fM | 400 fM | 800 fM | 1.6 pM | 3.2 pM |
| mean (N = 10) | 1027 | 1137 | 1100 | 1275 | 1392 | 1625 | 2347 | 4158 |
| SD | 86 | 147 | 104 | 131 | 84 | 208 | 229 | 451 |
| Threshold | 1884 | | | | | | | |
| Sample above threshold? | | nd | nd | nd | nd | nd | detected | detected |
| **N782 (1)** | no RNA | 10 fM | 20 fM | 40 fM | 80 fM | 160 fM | 320 fM | 640 fM |
| mean (N = 10) | 737 | 773 | 910 | 1156 | 938 | 1197 | 1468 | 2412 |
| SD | 84 | 100 | 132 | 237 | 89 | 99 | 162 | 121 |
| Threshold | 1579 | | | | | | | |
| Sample above threshold? | | nd | nd | nd | nd | nd | nd | detected |
| **N782 (2)** | no RNA | 10 fM | 20 fM | 40 fM | 80 fM | 160 fM | 320 fM | 640 fM |
| mean (N = 10) | 708 | 657 | 708 | 787 | 864 | 1045 | 1522 | 2328 |
| SD | 44 | 111 | 83 | 126 | 114 | 134 | 119 | 180 |
| Threshold | 1151 | | | | | | | |
| Sample above threshold? | | nd | nd | nd | nd | nd | detected | detected |
| **N782 (3)** | no RNA | 10 fM | 20 fM | 40 fM | 80 fM | 160 fM | 320 fM | 640 fM |
| mean (N = 10) | 780 | 808 | 765 | 900 | 981 | 1194 | 1719 | 2721 |
| SD | 94 | 124 | 108 | 95 | 139 | 106 | 147 | 173 |
| Threshold | 1719 | | | | | | | |
| Sample above threshold? | | nd | nd | nd | nd | nd | nd | detected |
| **N1169 (1)** | no RNA | 1 fM | 2 fM | 4 fM | 8 fM | 16 fM | 32 fM | 64 fM |
| mean (N = 10) | 559 | 641 | 753 | 984 | 1360 | 2399 | 5015 | 10250 |
| SD | 84 | 84 | 115 | 100 | 126 | 203 | 265 | 852 |
| Threshold | 1403 | | | | | | | |
| Sample above threshold? | | nd | nd | nd | nd | detected | detected | detected |
| **N1169 (2)** | no RNA | 500 aM | 1 fM | 2 fM | 4 fM | 8 fM | 16 fM | 32 fM |
| mean (N = 10) | 694 | 692 | 827 | 771 | 812 | 1130 | 1522 | 2632 |
| SD | 59 | 78 | 91 | 123 | 128 | 121 | 140 | 181 |
| Threshold | 1280 | | | | | | | |
| Sample above threshold? | | nd | nd | nd | nd | nd | detected | detected |
| **N1169 (3)** | no RNA | 500 aM | 1 fM | 2 fM | 4 fM | 8 fM | 16 fM | 32 fM |
| mean (N = 10) | 694 | 727 | 721 | 706 | 860 | 1005 | 1331 | 1975 |
| SD | 82 | 66 | 121 | 86 | 104 | 127 | 107 | 182 |
| Threshold | 1517 | | | | | | | |
| Sample above threshold? | | nd | nd | nd | nd | nd | nd | detected |
| **N209 (1)** | no RNA | Extract 6 | Extract 5 | Extract 4 | Extract 3 | Extract 2 | Extract 1 | SARS-Cov-2 |
| mean (N = 15) | 4432 | 4281 | 5717 | 20841 | 122614 | 164411 | 164551 | |
| SD | 385 | 381 | 345 | 1652 | 4955 | 3716 | 5617 | |
| Threshold | 8281 | | | | | | | |
| Sample above threshold? | | nd | nd | detected | detected | detected | detected | |
| **N209 (2)** | no RNA | Extract 1 | Extract 2 | Extract 3 | Extract 4 | Extract 5 | Extract 6 | SARS-Cov-2 |
| mean (N = 15) | 7537 | 170419 | 171694 | 132152 | 27056 | 12308 | 8939 | |
| SD | 782 | 1788 | 1979 | 8468 | 4743 | 3676 | 2323 | |
| Threshold | 15360 | | | | | | | |
| Sample above threshold? | | detected | detected | detected | detected | nd | nd | |

The tested VmeCmr complexes are indicated by their crRNA target. The threshold was defined as the reference mean plus 10 reference standard deviations. If the sample mean was above the threshold, the sample was deemed “detected”, and “nd” (not detected) if it was below.

**Table S2. Ct values from RT-qPCR analysis.**

| 1 aM [a] | 10 aM [a] | 100 aM [a] | 1 fM [a] | 10 fM [a] | 100 fM [a] | 1 pM [a] |
|---|---|---|---|---|---|---|
| 36.7 | 31.29 | 28.14 | 24.78 | 21.67 | 18.28 | 15.06 |
| No Ct | 31.53 | 28.07 | 24.92 | 21.57 | 18.41 | 15.36 |
| 38.71 | 31.76 | 28.13 | 25.01 | 21.51 | 18.58 | 15.23 |
| | | | | | | **NRT** |
| | | | | | | 33.74 |
| | | | | | | 33.53 |
| | | | | | | 34.08 |

| NTC | Extract 1 [b] | Extract 2 [b] | Extract 3 [b] | Extract 4 [b] |
|---|---|---|---|---|
| 38.34 | 14.74 | 18.24 | 21.90 | 25.15 |
| No Ct | 14.96 | 18.33 | 22.00 | 25.35 |
| No Ct | 14.75 | 18.42 | 21.83 | 25.29 |
| | | **NRT** | **NRT** | **NRT** |
| | | No Ct | No Ct | No Ct |
| | | No Ct | No Ct | No Ct |
| | | No Ct | No Ct | No Ct |

a) SARS-CoV-2 N gene transcript
b) SARS-CoV-2 genomic RNA extract
NTC: no template control; NRT: no reverse transcriptase control
